# A highly homogeneous expansion microscopy polymer composed of tetrahedron-like monomers

**DOI:** 10.1101/814111

**Authors:** Ruixuan Gao, Chih-Chieh (Jay) Yu, Linyi Gao, Kiryl D Piatkevich, Rachael L Neve, Srigokul Upadhyayula, Edward S Boyden

**Affiliations:** McGovern Institute for Brain Research, MIT, Cambridge, MA, USA; MIT Media Lab, MIT, Cambridge, MA, USA; Janelia Research Campus, Howard Hughes Medical Institute, Ashburn, VA, USA; Department of Biological Engineering, MIT, MA, USA; Broad Institute, MIT, Cambridge, MA, USA; Department of Neurology, Massachusetts General Hospital, Cambridge, MA, USA; Department of Cell Biology, Harvard Medical School, Boston, MA, USA; Program in Cellular and Molecular Medicine, Boston Children’s Hospital, Boston, MA, USA; Department of Pediatrics, Harvard Medical School, Boston, MA, USA; Advanced Bioimaging Center, University of California at Berkley, Berkeley, CA, USA; Department of Molecular and Cell Biology, University of California at Berkeley, Berkeley, CA, USA; MIT Center for Neurobiological Engineering, MIT, Cambridge, MA, USA; Department of Brain and Cognitive Sciences, MIT, Cambridge, MA, USA; Koch Institute, MIT, Cambridge, MA, USA

## Abstract

Expansion microscopy (ExM) physically magnifies biological specimens to enable nanoscale-resolution imaging on conventional microscopes. Current ExM methods permeate biological specimens with free radical-polymerized polyacrylate hydrogels, whose network structure limits the microscopy resolution enabled by ExM. Here we report that ExM is possible using hydrogels with more homogeneous network structure, assembled via non-radical terminal linking of monomers of tetrahedral shape. As with earlier forms of ExM, such “tetra-gel”-embedded specimens can be iteratively expanded for greater physical magnification. Iterative tetra-gel expansion of herpes simplex virus type 1 (HSV-1) virions by ~10x in linear dimension results in a viral envelope deviation from sphericity of 9.2 nm, rather than the 14.3 nm enabled by free radical-polymerized hydrogels used in earlier versions of ExM. Thus, tetra-gel polymer chemistry may support new forms of ExM imaging that introduce fewer spatial errors than earlier versions, and raise the question of whether single biomolecule precision may be achievable.

## MAIN

Expansion microscopy (ExM) is in increasingly widespread use because it enables, via physical magnification of biological specimens, nanoscale imaging on conventional microscopes^1–3^. In ExM, biological specimens are first embedded in a swellable hydrogel network, chiefly composed of crosslinked sodium polyacrylate. During this embedding, biomolecules and/or fluorescent tags are covalently anchored to the hydrogel network, so that their relative organization is preserved after the specimen is enzymatically or chemically softened, and the hydrogel expanded upon immersion in water (typically ~4x-5x in linear dimension). Protocols using off-the-shelf-chemicals^4^ have helped ExM find utility in a wide variety of contexts, ranging from the mapping of ribosome components and RNAs in presynaptic compartments^5^, to the analysis of circadian rhythm neural circuitry in the *Drosophila* brain^6^, to the analysis of kidney disease and breast cancer in human biopsies^7^, with new results appearing continuously (see review^3^). Variants of ExM have been developed that achieve nanoscale localization of proteins and RNAs in preserved cells and tissues on diffraction-limited microscopes^4,7–13^. Additionally, new strategies have been introduced to expand specimens ~10x-20x in linear dimension, by applying the expansion process over and over (iterative expansion microscopy or iExM)^14^ or by using superabsorbent hydrogels (X10 expansion microscopy)^15^. However, all ExM variants to date form the hydrogel mesh via free-radical polymerization, a process that results in nanoscale structural heterogeneity^16–20^. This raises the question of whether another polymer, or polymerization process, might result in better structural heterogeneity, and thus less spatial error during the expansion process.

Before proposing a strategy, we describe two earlier ExM protocols in detail to provide context, since we aim to improve specific steps of these processes. One popular version of ExM is so-called protein-retention expansion microscopy (proExM)^8^. In one variant of proExM, fixed biological specimens that have been labeled with fluorophore-bearing antibodies are exposed to a small molecule (abbreviated as AcX) that equips the primary amine groups on antibodies and proteins with a polymerizable acryloyl group. The specimen is then immersed in a solution containing sodium acrylate monomer (with acrylamide co-monomer), as well as the crosslinker N,N-methylenebisacrylamide, and a dense network of crosslinked sodium polyacrylate hydrogel is then formed, permeating throughout the specimen, through free-radical polymerization. During this process, the antibodies and proteins are covalently anchored to the hydrogel polymer network via the AcX linker. Finally, an enzymatic digestion step using proteinase K cleaves most of the proteins, largely sparing the antibody-targeted fluorophores, and allowing for expansion of the polymer-specimen composite in water. As a second example of an ExM protocol, iterative expansion microscopy (iExM)^14^, fixed specimens are first labeled with primary antibodies and then DNA oligo-conjugated secondary antibodies. Next, oligos bearing a gel-anchorable moiety are hybridized to the secondary antibody-conjugated DNA oligos. Then a first hydrogel is formed as above, but with a cleavable cross-linker, which anchors the gel-anchorable DNA oligos to the hydrogel at the locations of the immunostained proteins; this gel is then expanded as described above for proExM. Next, another round of hybridization of DNA oligos complementary to those anchored to the first hydrogel, and again bearing a gel-anchorable moiety, is performed. Then, a second hydrogel is formed throughout the expanded first hydrogel, so that the new DNA oligos are anchored to the second hydrogel at the sites of immunostained proteins. Finally, the first hydrogel is cleaved, allowing the second hydrogel to expand upon immersion in water. The protein locations can be imaged after the polymer-anchored oligos are labeled by applying complementary oligos, potentially equipped with branched DNA for amplification purposes, bearing fluorescent dyes. The oligos, in short, provide a molecular mechanism (based on complementary strand hybridization) for signal transfer between the first and second hydrogels, as well as a scaffold for amplification and fluorescence readout.

In both cases, the free-radical polymerization process that forms the sodium polyacrylate hydrogel has nanoscale structural heterogeneity. A structural study using small-angle X-ray scattering (SAXS) found that the mean size of local cross-linker density variations (Fig. 1a, “1”) can amount to 15-25 nm for polyacrylamide gels, depending on the gels’ monomer and cross-linker concentrations^16^. Moreover, topological defects of the polymer chains, such as dangling ends (Fig. 1a, “2”) and loops (Fig. 1a, “3”), are known to introduce deviations from uniform polymer network meshes at the 1-10 nm length scale^19,20^. So far, no studies have directly and precisely assessed whether and how the structural heterogeneity of the hydrogel matrix relates to the microscopy resolution enabled by ExM, and therefore it remains unclear at this point how much the aforementioned monomer/cross-linker density variations and topological defects would affect the effective resolution of ExM. In our recent study of iExM, described in the previous paragraph, we found that the process of expanding cells ~20x in two rounds of gel formation and expansion, resulted in ~9-13 nm of local distortion for the expansion of tags targeted via antibodies to tubulin^14^.

**Figure 1.**
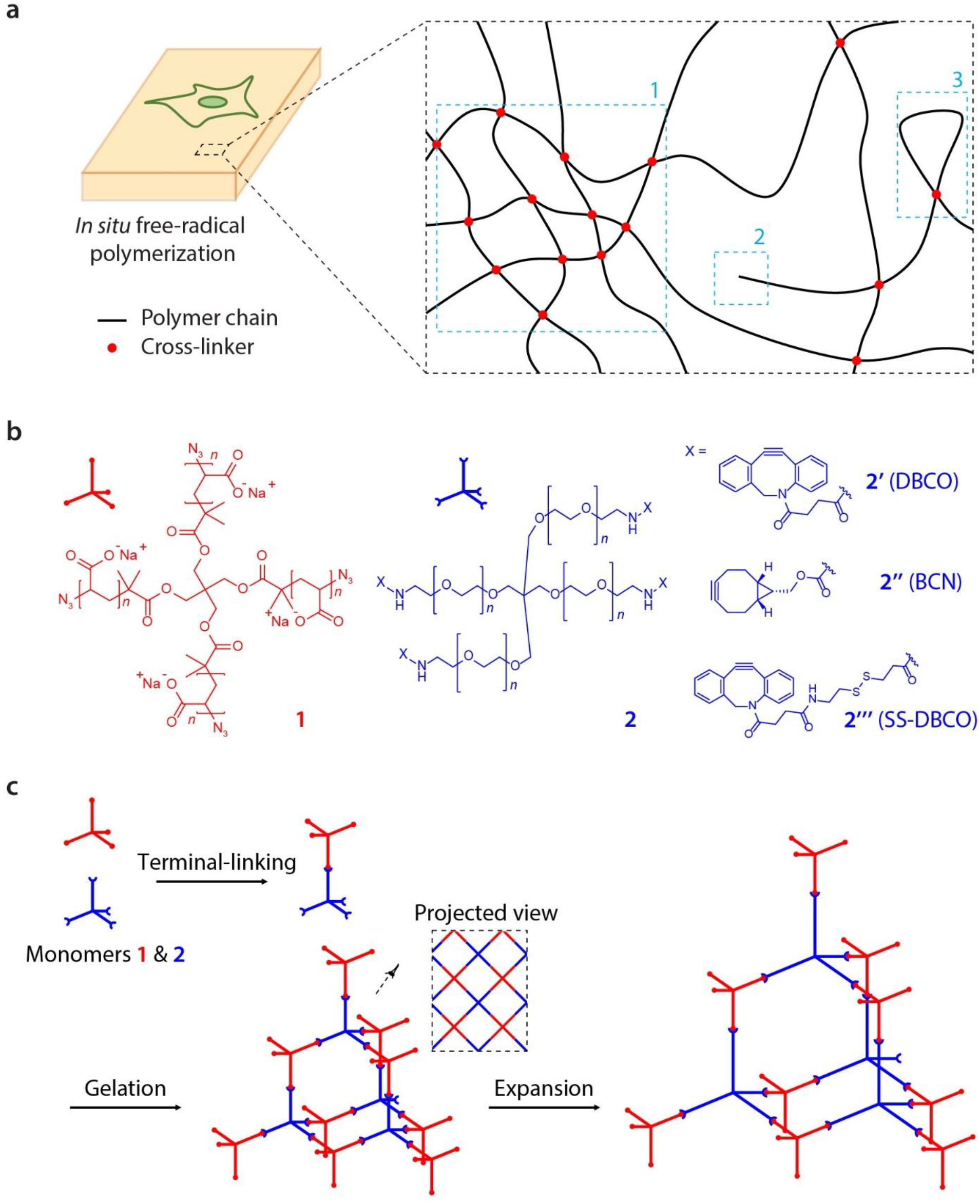
Design and synthesis of tetra-gels (TG) for expansion microscopy. **a**, Cell/tissue-hydrogel composites formed by *in situ* free-radical polymerization are known to have structural inhomogeneities in the range of tens of nanometers due to (1) local fluctuations of monomer and cross-linking densities, (2) dangling ends, and (3) loops formed within the polymer network. **b**, Design of TG monomers **1** and **2** with tetrahedral symmetry and reactive terminal groups. Specific monomer **2** terminal groups (**2’**, **2’’** and **2’’’**) enable, for example, control of the reaction rate between the monomers and addition of functionalities to the polymer network. **c**, Formation and expansion of TG via click-chemistry-based terminal-linking of monomers **1** and **2**. Inset, projected view of the TG polymer network.

In the field of polymer chemistry, many attempts have been made to obtain an ideal hydrogel matrix, with a more homogenous polymer network than obtainable with free-radical polymerization^19–24^. One study utilizing terminal linking of two kinds of tetrahedral polyethylene glycol (PEG) monomers to form a diamond lattice-like polymer network resulted in hydrogels with extremely high structural homogeneity^19,20,24^. Small-angle neutron scattering (SANS) and static light scattering (SLS) studies showed that the thus-synthesized hydrogel, called a tetra-PEG gel, was structurally proximate to an ideal polymer network, nearly void of structural defects such as loops, dangling chains, and trapped entanglements at both the as-synthesized state and the slightly swollen state (<1.5x linear expansion in water)^25,26^. Recent studies have demonstrated that by swapping the terminal functional groups or the tetra-arm backbones with other chemical moieties, the tetra-PEG gel structure can be exploited to create hydrogels with versatile chemical and mechanical properties, such as click-chemistry-based polyelectrolyte hydrogels^27^ and highly compressible and stretchable hydrogels^28^. In the tetra-PEG design, the polymer chain lengths and the cross-linking densities are highly uniform owing to the consistent monomer size and the complementary, mutually-limiting polymerization mechanism. Moreover, topological defects of the polymer network, such as loops and dangling ends, are largely removed by design, due to the specific and stochiometric terminal linking.

We here report a class of swellable hydrogels based on the tetra-PEG structure, designed to have structural heterogeneity better than that of the radically polymerized hydrogel matrices used in previous ExM protocols. Formed by click-chemistry-based, non-radical linking of two complementary, tetrahedral monomers comprising backbones of polyacrylate and PEG respectively, and called tetra-gel (TG), the resulting optically transparent hydrogels swell ~3.0-3.5-fold linearly in deionized water. By infusing preserved cells or tissues with a small molecule linker such as NHS-azide, which adds a click-reactive azide group to primary amines of biomolecules such as proteins, we show that proExM can be performed with the TG polymer. Moreover, introduction of a cleavable moiety to one of the monomers allows the formed TG to be cleaved into individual monomers after being expanded and then re-embedded in a second hydrogel, rendering TG compatible with the iExM process^14^. We investigated whether TG-based iExM improved resolution over our earlier form of iExM. Given that resolution and spatial error of iExM is in significant part limited by the size of the antibodies administered pre-expansion^14^, for the purposes of evaluating TG-based iExM chemistry, we implemented a direct-labeling strategy to directly conjugate DNA oligos nonspecifically to protein targets. Using herpes simplex virus type 1 (HSV-1) virions as a testbed, we found that ~10-fold expanded HSV-1 virions exhibited envelope proteins with significantly smaller deviation from a circular shape at the virion midplane, compared to the classical sodium polyacrylate/acrylamide gel (PAAG)-expanded virions [TG: 9.2 nm (median), PAAG: 14.3 nm (median); *p* < 10^−20^, 2-sided Wilcoxon rank sum test; TG: 352, PAAG: 330 virion particles; all virions from the same single batch of live HSV-1 preparation]. Although new labeling strategies will be required for TG-based iExM to exhibit this enhanced precision in visualizing specific biomolecule types, the chemistry discoveries here reported show that structurally homogeneous expansion microscopy matrices are possible, and point the way to new ways to generalize, improve, and extend the expansion microscopy toolbox.

## RESULTS

### Design of the tetra-gel (TG) monomers and polymerization chemistry

Free-radical polymerization introduces nanoscale structural heterogeneity into hydrogel networks at the length scale of tens of nanometers^16–19^, due to (1) local fluctuations of the monomer and cross-linker densities, as well as topological defects of the polymer chains, such as (2) dangling chain ends and (3) loops (Fig. 1a). We designed and synthesized tetrahedral monomers closely related to those known to form the homogenous tetra-PEG structure via non-radical terminal-linking (Fig. 1b, **Supplementary Fig. 1**)^24–26^. In our hydrogel design, one of the monomers (monomer **1**) has a tetra-arm polyacrylate backbone with a clickable terminal group (azide), and the other (monomer **2**) has a tetra-arm PEG backbone with a complementary terminal group (alkyne). The alkyne is synthetically switchable to tune the reactivity of the terminal linking and/or to add functionalities to the hydrogel network (Fig 1b, monomers **2’, 2’’, 2’’’**). We designed these monomers so that they had a comparable molecular weight of ~10-20k Da and an arm length of ~3-6 nm at the gelation step (at an ionic strength of ~0.150M). In more detail: in the solution phase, Monomer **1** has four negatively charged polyacrylate arms (n = ~21 monomer units long), each arm of which is estimated to extend ~4.0-5.7 nm, based on the previously characterized persistence length of ≥ 4.0 nm^29^ and the fully stretched length of ~5.7 nm. Monomer **2** has four uncharged PEG arms (n = either ~57 or ~114 monomer units long, depending on which design is utilized), each of which has a fully stretched length of ~21.7 or ~43.3 nm, respectively, which greatly exceeds its persistence length (0.38 nm)^30^. The PEG arm can thus be modeled as a freely jointed chain in solution^31,32^, whose root-mean-square end-to-end distance (i.e., the arm length) can be calculated as either ~2.9 or ~4.1 nm, respectively. For monomer **2** synthesis, the n = ~57 backbone was used for variants **2’** and **2’’** and the n = ~114 backbone was used for variant **2’’’**; the increased number of PEG units in **2’’’** enhanced its solubility to compensate for the hydrophobic nature of the SS-DBCO moiety. In this design, when monomer **1** and a selected monomer **2** are mixed, a hydrogel would form via click-chemistry-based terminal linking, similar to tetra-PEG gel formation (Fig. 1c, left). Then, in deionized water or aqueous buffers with an ionic strength smaller than ~0.05M (concentration estimated from the Debye-Huckel theory for the electrostatic potential energy in electrolyte solutions)^33^, the hydrogel would swell due to the reduced electrostatic screening of salt ions between the mutually repelling monomer **1** units (~84 negative charges per monomer), which in turn would elongate the originally unstructured PEG arms of the interconnecting monomer **2** (Fig. 1c, right).

Of the monomer **2** variants in Fig. 1b, the bicyclononyne (BCN) version (monomer **2’’**) was designed to support expansion of thicker tissues than the standard dibenzocyclooctyne (DBCO) version (monomer **2’**), because its slower click-reaction kinetics (by ~55%)^34^ would provide additional time for the monomers to diffuse, and equilibrate in density, throughout the tissue during the pre-gelation incubation step. Moreover, the disulfide dibenzocyclooctyne (SS-DBCO) version (monomer **2’’’**) would enable post-expansion cleavage of the polymer network into individual monomers after being reembedded in a second hydrogel, rendering it compatible with iterative expansion^14^.

### TG expansion of cells and tissues

We mixed monomers **1** and **2**, targeting a stoichiometric ratio of 1:1, and casted the gelling solution into a circular mold. The molar ratio of monomer **1** to **2** in the initial recipe could deviate slightly from 1:1 depending on the fraction of monomer **1** whose t-butyl group was removed in the final step of monomer **1** synthesis (**Supplementary Fig. 1**, bottom row), which determines the exact molecular weight of monomer **1**. Nonetheless, the mutually-limiting click polymerization mechanism of TG formation should assure a 1:1 stochiometric incorporation of monomers **1** and **2** during gel formation, even if concentrations are slightly different. After 1-2 hours of incubation at 37°C, we found that the gelling solution solidified into an optically transparent and mechanically elastic hydrogel (Fig. 2a, left). Similar to the radically formed PAAG gel and other types of hydrogels used in ExM and tissue clearing methods^7–13^, the TG swelled after salt elution in water (~3-fold in linear dimension) (Fig. 2a, right). We anchored a small amount of fluorescent dye to the TG polymer network, and imaged fluorescently-labeled gels before and after swelling (Fig. 2b). We found that the linear expansion factor of TG was ~3.0-3.5x.

**Figure 2.**
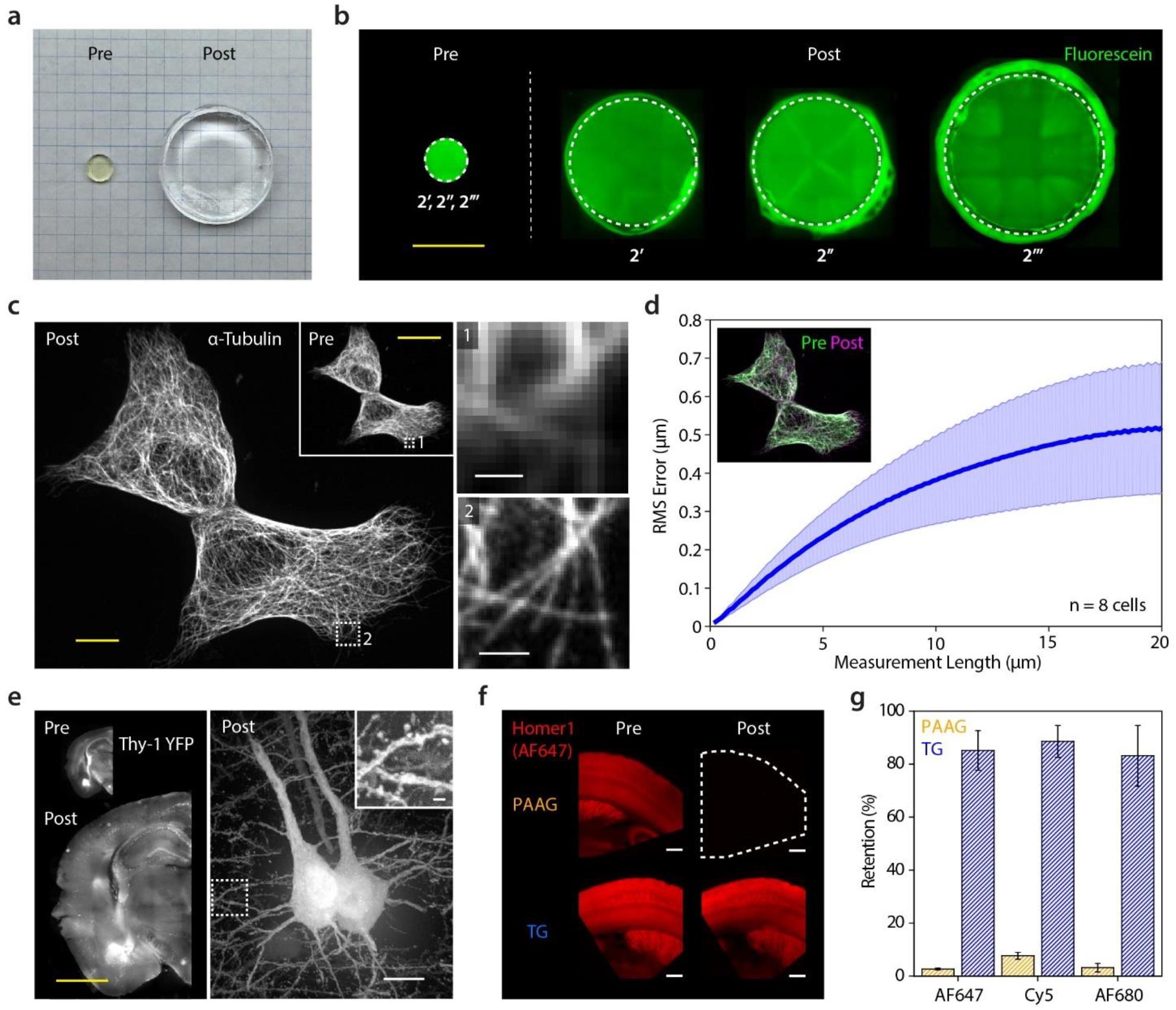
TG-mediated expansion of cells and tissues. **a**, Image of TG (using monomer **2’’**) as synthesized (left, pre-expansion) and after swelling in deionized water (right, post-expansion). The two gels were cast in circular molds with identical dimensions. Grid size, 5 mm. **b**, TGs labeled with fluorescein in pre-(left, **2’**; same sizes and shapes when **2’’** and **2’’’** were used) and post-expansion (right, **2’**, **2’’** and **2’’’**) states. Irregular boundaries on the post-expansion images reflect the meniscus of water used to expand the gels. Scale bar, 5 mm. **c**, Left, HEK293 cells with α-tubulin immunostaining in pre-(inset) and post-expansion states. Expansion factor, 2.85x. Scale bars, 20 µm. Right, magnified views of the boxed regions 1 (top) and 2 (bottom). Scale bars, 1 µm [bottom, 2.85 µm; here and after, unless otherwise noted, scale bar sizes are given at pre-expansion scale (i.e., biological scale) with the corresponding post-expansion size (i.e., physical size) indicated in parentheses]. **d**, Root-mean-square (RMS) error curve for HEK293 cell expansion (blue line, mean; shaded, standard deviation; n = 8 cells from one culture). Inset, non-rigidly registered and overlaid pre-(green) and post-expansion (magenta) images used for the RMS error analysis. **e**, Pre-(left top) and post-expansion (left bottom and right) Thy1-YFP mouse brain slices. Expansion factor, 3.00x. Scale bars, 5 mm (left) and 10 µm (right, 30 µm). The gelled brain slice on the left bottom panel was immunostained against YFP after the proteolysis step to enhance fluorescence. Inset, magnified view of the boxed region. Scale bar, 1 µm (3 µm). **f**, Pre-expansion (left column) and post-expansion (right column) Thy1-YFP mouse brain slices immunostained with Homer1 primary antibody and Alexa Fluor 647 (AF647)-conjugated secondary antibody, using sodium polyacrylate/acrylamide gel (PAAG, top row) and TG (bottom row). Expansion factors, 3.93x (PAAG) and 2.72x (TG). Scale bars, 300 µm (top right, 1.18 mm; bottom right, 815 µm). **g**, Fluorescence retention of AF647, Cy5, and AF680 with PG and PAAG in brain slices processed as in **f** (bar height, mean; error bar, standard error of the mean; n = 3 brain slices from one mouse).

We implemented proExM using TG. As mentioned earlier, a small-molecule linker, AcX, is reacted to primary amines on proteins (either endogenous, or exogenously applied such as fluorescent dye-conjugated antibodies) in the biological specimen, allowing them to be covalently anchored to, and physically separated from one another by, the swelling polymer network^4,8^. We infused antibody-stained cells and tissue slices with a small-molecule linker, NHS-azide, so that the primary amines of the proteins and antibodies could be covalently bound to the polymer network through azide-based click-chemistry (Fig. 1b). We then formed the TG *in situ* before mechanically homogenizing the cell/tissue-TG composites by proteinase K proteolysis, as in the proExM process^4,8^. We note that NHS-azide can be potentially replaced by other small molecules, as long as they covalently bind to the target biomolecules as well as either the terminal or side functional groups of the TG polymer network.

For TG-based expansion of cultured cells, we embedded HEK cells immunostained against microtubules in the DBCO version (monomer **2’**) of TG. After swelling in deionized water, the HEK cell-TG composite expanded ~2.9-fold, resulting in more sharply resolved microtubules than before expansion, on the same diffraction-limited confocal microscope (Fig. 2c). We aligned pre- and post-expansion confocal microscopy images of microtubules via non-rigid registration, and quantified the amount of distortion in the expansion process at a macro length scale (Fig. 2d), finding ~2.5% error over measurement lengths of 20 µm, comparable to the macro distortion of earlier PAAG-based proExM^1,8^. We further embedded mouse brain slices expressing yellow fluorescent protein (YFP) in the slower-to-react BCN version (monomer **2’’**) of TG, and immunostained against YFP (whose protease resistance we utilized in the earlier PAAG-based proExM protocol)^8^ after proteinase K proteolysis, to enhance fluorescence (Fig. 2e). After swelling, the mouse brain slice-TG composite expanded ~3.0-fold and showed enhanced detail in subcellular structures such as dendritic spines, as observed before with PAAG-based proExM^8^.

Since TG forms via non-radical polymerization, TG-based expansion could potentially preserve molecules that are destroyed or modified during radical polymerization. To test this, we measured fluorescence retention of small-molecule dyes that are known to be susceptible to free-radical polymerization in PAAG-based proExM (Fig. 2f). We found that the fluorescence of cyanine dyes, including Alexa Fluor 647 (AF647), Cy5, and AF 680, was nearly lost (<10% retention, Fig. 2g) after PAAG-based proExM, as observed before^8^, but was largely preserved (>80% retention) after TG-based expansion (Fig. 2f, 2g). TG-based ExM is thus compatible with pre-expansion staining with fluorescent dyes not compatible with PAAG-based ExM^35^.

### TG-based iterative expansion

The SS-DBCO version of TG (using monomer **2’’’**) allows the TG polymer network to be cleaved at every node that connects two neighboring monomers (Fig. 3a). Such cleavability makes TG well suited as the first-round hydrogel for iterative expansion^14^. To verify TG’s cleavability and compatibility with iExM, we applied the previously described principle of iterative expansion to TG-expanded HeLa cells (Fig. 3b). In this process, HeLa cells were stained with primary antibodies against tubulin followed by DNA oligo-conjugated secondary antibodies, embedded in the SS-DBCO (monomer **2’’’**) version of TG, proteolytically digested with proteinase K, expanded by deionized water, and re-embedded in a PAAG for the 2nd around of expansion and imaging. We used PAAG for the second round of expansion, reasoning that the spatial errors introduced by PAAG-based expansion in the second round would be negligible. This is because the spatial errors introduced by PAAG-based expansion in the second round, when considered in biological units (i.e., in terms of the relative spacing of biomolecules with respect to each other), are effectively divided by the expansion factor of the first round, and thus would be ~3x smaller than those that would be introduced by using PAAG in the first round of iterative expansion. After cleaving the disulfide bonds of the TG with tris(2-carboxyethyl)phosphine (TCEP) and adding deionized water, the cells expanded by ~16-fold. Consistent with previously reported structures in BS-C-1 cells expanded by PAAG-based iterative expansion^14^, we could detect the hollow of microtubules after TG-based iterative expansion (Fig. 3c). Quantitative analysis of the peak-to-peak distance between the microtubule sidewalls shows an average distance of ~65.3 +/− 15.7 nm (mean +/− std. dev.; n = 336 microtubule segments of 200 nm length, from 10 cells in one culture), comparable to that seen previously with iExM of BS-C-1 cells using PAAG gels (Fig. 3d). Multiple DNA oligo-conjugated antibodies bearing different oligo sequences could be used at once, e.g. to label tubulin and clathrin in HeLa cells (Fig. 3e)^14^.

**Figure 3.**
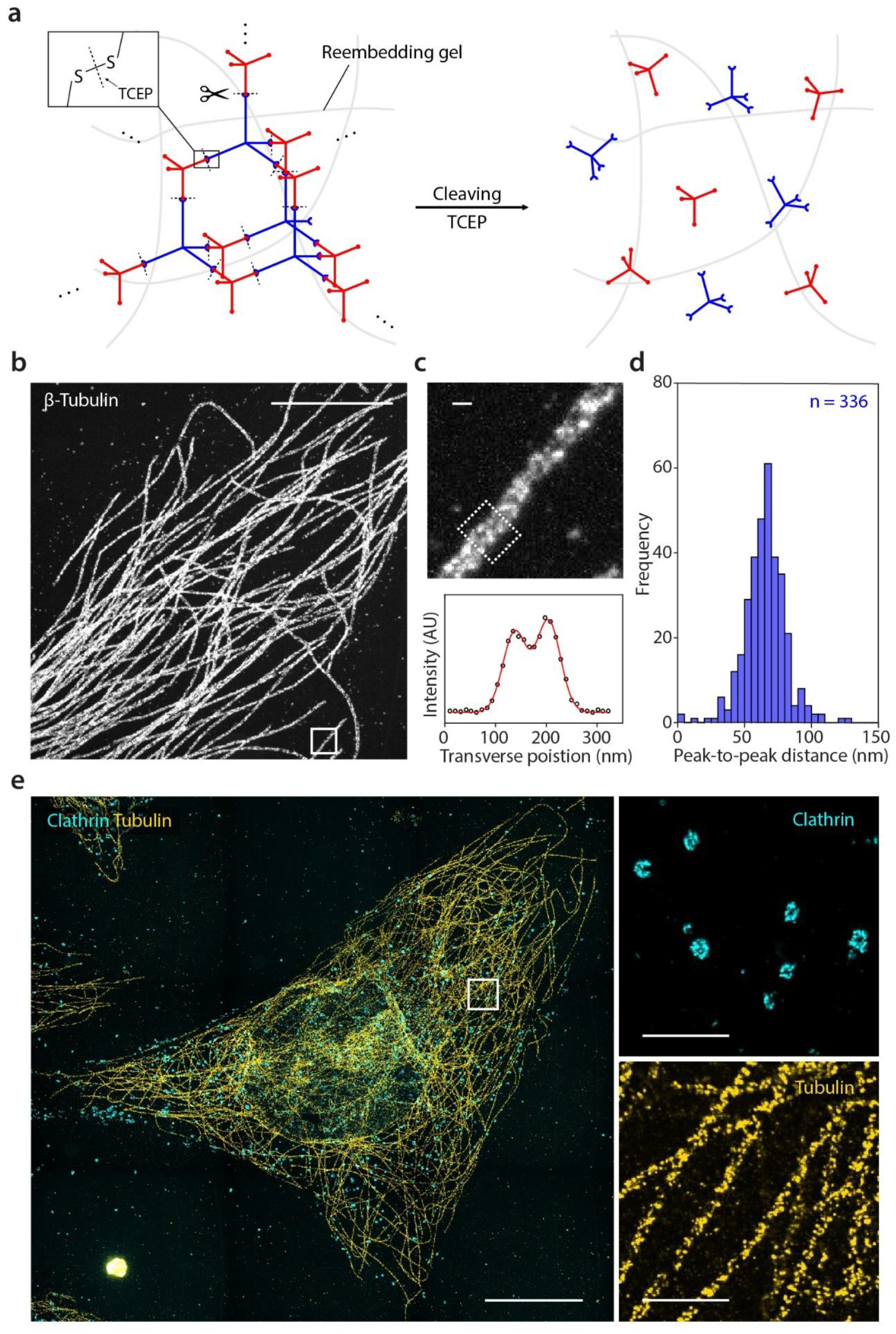
TG-based iterative expansion. **a**, Monomeric cleaving of TG (monomer **2’’’**) after reembedding in a second hydrogel. A reducing agent, tris(2-carboxyethyl)phosphine) (TCEP), was applied to cleave the disulfide bonds in the TG polymer network. **b**, HeLa cell with β-tubulin immunostaining, expanded by cleavable TG-based 2-round iterative expansion. Expansion factor, 15.6x. Scale bar, 5 µm (78.2 µm). **c**, Top, magnified view of the boxed region in **b**. Scale bar, 100 nm (1.56 µm). Bottom, transverse line intensity profile of the microtubule in a single xy-plane in the dotted box (circle) and the fitted sum of two Gaussians (red line). The line intensity profile was averaged in the direction parallel to the microtubule axis over a length of 200 nm. **d**, Histogram of peak-to-peak distance between microtubule sidewalls in HeLa cells (n = 336 segments of a length of 200 nm from 10 cells in one culture). **e**, Left, HeLa cell with two-color labeling of clathrin coated pits/vesicles and microtubules, expanded by TG-based 2-round iterative expansion. Expansion factor, 15.6x. Scale bar, 10 µm (156 µm). Right, magnified view of the boxed region for each color channel. Scale bars, 1 µm (15.6 µm).

### Expansion of virions and spatial arrangements of envelope proteins

To assess the spatial errors introduced by TG-based iExM, we imaged the nanoscale arrangement of herpes simplex virus type 1 (HSV-1) virion envelope proteins. In iExM, primary and secondary antibodies, and the conjugated DNA oligos used in the process of iExM, have been estimated to add ~21 nm of spatial error to the localization of epitopes^14^, potentially confounding the assessment of any fine-scale improvements of TG over PAAG forms of iExM. In order to avoid this limitation, wherein pre-expansion administration of antibodies against epitopes means that antibody size is a major contributor to the spatial errors introduced by iExM^14^, we developed a direct, nonspecific labeling strategy that targets primary amines of envelope proteins. For the labeling, we directly conjugated 22-bp oligonucleotides to the envelope proteins on HSV-1 virions via a hydrazone-formation-based DNA-to-peptide conjugation (Fig. 4a). This direct labeling protocol reduces the label size from ~21 nm to that of a single DNA oligo (~7 nm)^14^. We are careful to not expose viruses to any buffers containing detergents (e.g. Triton), for all steps from fixation to gelation, aiming to preserve the virus’s lipid bilayer envelope integrity, so that the highly negatively charged 22-mer DNA oligo will unlikely be able to cross the membrane, and thus conferring specificity of the labeling to the accessible envelope proteins on the outside of the membrane. Of course, the nontargeted nature of this labeling scheme means that the chemistry discoveries here reported will need further innovation in order for specific proteins to be labeled, e.g. by post-expansion delivery of antibodies (which has been performed for single rounds of expansion^8,11^ but has not yet been reported for the iterative expansion case), which would result in antibody-contributed spatial errors that are effectively divided by the expansion factor. Given that our goal for the current manuscript was to explore the foundational chemistry of expansion microscopy hydrogels, we decided to focus on the fundamental chemistry of TG-based iExM, as opposed to developing a practical protocol for biologists interested in specific proteins; in the future, such a protocol could build upon the chemistries and discoveries of the current chemistry study.

**Figure 4.**
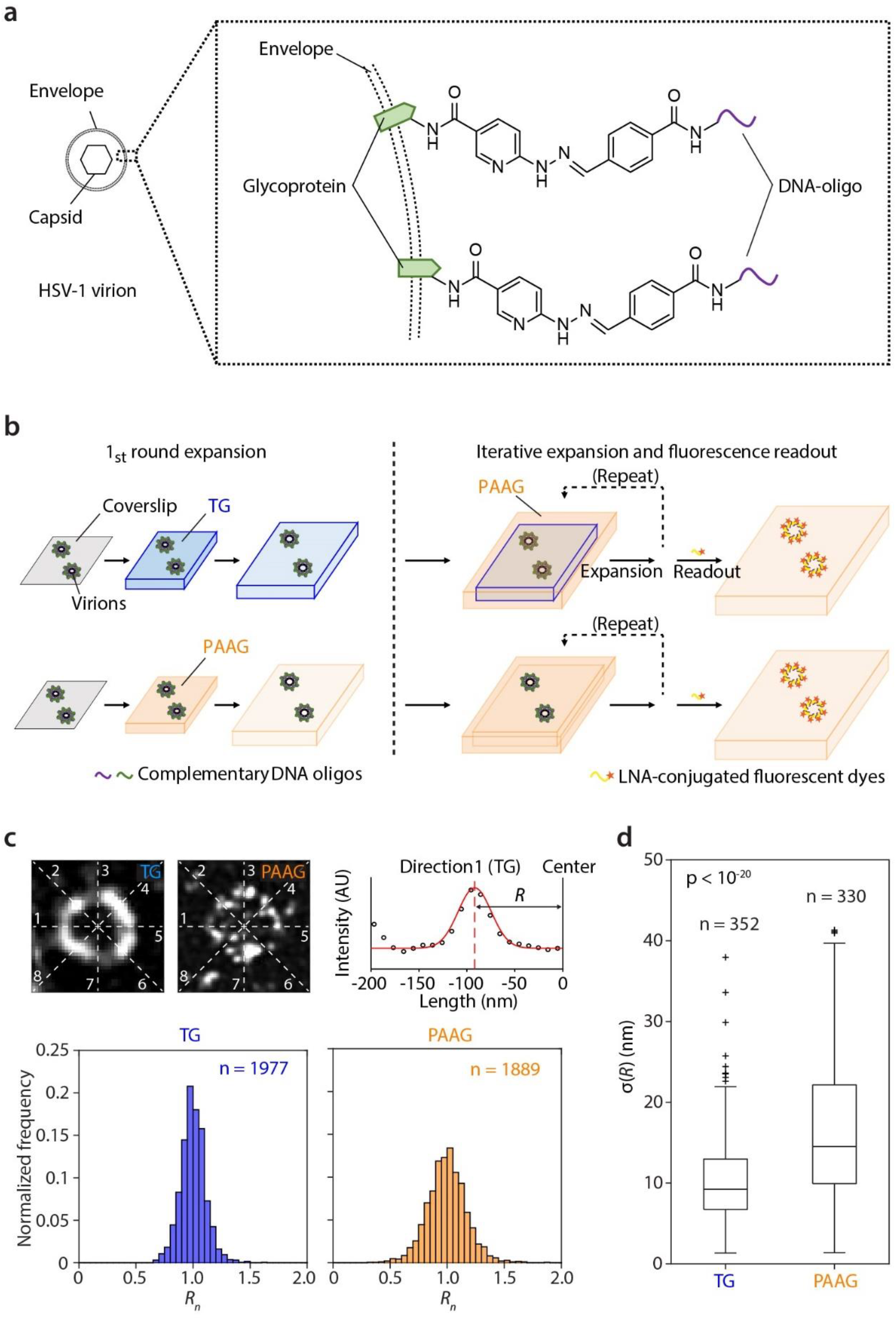
Spatial errors introduced by TG-based vs. classical PAAG-based iterative expansion microscopy. **a**, Short DNA-oligos (22 bp) were covalently conjugated to the envelope proteins of herpes simplex virus type 1 (HSV-1) virions via hydrazone formation, which allows labeling transfer across multiple hydrogels, amplification based on branched DNA, and fluorescence readout based on hybridization of fluorescent oligos. **b**, Schematic illustration of TG-(top) and PAAG-based (bottom) iterative expansion of HSV-1 virions with the direct oligo-conjugation of **a**. PAAG-based expansion was used for all expansion rounds after the first round, reasoning that most of the error of an iterative expansion protocol is introduced in the first round of expansion (see text for details). **c**, Top left and middle, representative single xy-plane images of HSV-1 virions expanded by TG- and PAAG-based 2-round iterative expansion. White lines indicate the 8 directions along which the virion particle’s radii (*R*) were measured. Size of both fields of view at biological scale, 400 nm. Top right, representative line intensity profile (circles) along a single direction (Direction1) of the TG-expanded virion and the fitted Gaussian (red line). Distance from the center of the Gaussian to the center of the virion was defined as *R* along that direction. Bottom, histogram of the particle-mean-normalized *R* (*R*_*n*_) for all the measured line profiles [TG, n = 1977; PAAG, n =1889 profiles; from 352 (TG) and 330 (PAAG) virion particles from the same single batch of live HSV-1 preparation]. **d**, Box plots [plus sign, outliers (values above 75^th^ percentile + 1.5 × interquartile range, or below 25^th^ percentile – 1.5 × interquartile range); ends of whiskers, maximum and minimum values of the distribution after outliers are excluded; upper line of box, 75^th^ percentile; middle line of box, 50^th^ percentile; lower line of box, 25^th^ percentile] of the standard deviation (*σ*) of HSV-1 virion radii (*R*) within individual virion particles for TG- and PAAG-based iterative expansion (*p* < 10^−20^, 2-sided Wilcoxon rank sum test; TG, n = 352; PAAG, n = 330 virion particles; virion particles from the same single batch of live HSV-1 preparation).

HSV-1 virions have a well-defined envelope protein layer that has been characterized by high-resolution imaging methods such as electron microscopy (EM)^36–38^, electron tomography (ET)^39,40^, and super-resolution microscopy^41^; are mechanically robust and tightly crosslinked after fixation^42^; have appropriate length scale (170-190 nm)^41^, and features in the tens of nanometers, to characterize the local structure of the TG hydrogel; and are compatible with the direct-labeling strategy here utilized. We directly labeled envelope proteins and expanded HSV-1 virions with the SS-DBCO (monomer **2’’’**) version of TG, comparing to PAAG, and then the gelled specimens were expanded a second time (in both cases using PAAG, as described above) to ~10-fold (Fig. 4b). This protocol was compatible with dual-color imaging of HSV-1 virion envelope proteins and DNA (**Supplementary Fig. 2**), as well as imaging of envelope proteins from other types of virions, such as vesicular stomatitis virus (VSV) virions (**Supplementary Fig. 3**).

We compared the spatial arrangement of proteins of the envelope layers of HSV-1 virions expanded by 2-round iterative expansion with either TG vs. PAAG serving as the first-round expandable gel (**Supplementary Fig. 4a**). The expansion factor for TG-based iExM of HSV-1 was ~10.3-13.3, and for PAAG-based iExM, it was ~15.3-18.7. From single-particle averaged virion images, we found that the envelope protein profiles of TG-expanded virions appeared significantly sharper than those of PAAG-expanded ones, suggesting reduced local spatial error for the former (**Supplementary Fig. 4b**). To quantify this spatial error, we measured the standard deviation (*σ*) of the virus radius within the mid-plane of individual HSV-1 virions (Fig. 4c). We computed this in biological units, normalizing by the expansion factor, to compensate for the different expansion factors for TG-based and PAAG-based iExM protocols. Previous cryo-ET studies^39,40^ found that the envelope of HSV-1 virions is a smooth, continuous, and near-spherical layer. We found that TG-based iExM-expanded virions [*σ* = 9.2 nm (median); n = 352 virion particles; virion particles from a single batch of live HSV-1 preparation] had significantly smaller median *σ* value compared to PAAG-expanded virions [*σ* = 14.3 nm (median); n = 330 virion particles; virion particles from a single batch of live HSV-1 preparation, same batch as TG] (*p* < 10^−20^, 2-sided Wilcoxon rank sum test) (Fig. 4d), suggesting a reduced spatial error mediated by TG-based vs. PAAG-based iExM. Even more resolution, potentially approaching that of individual proteins or protein clusters within HSV-1 virions, may be possible by expanding ~40x via 3 rounds of iterative expansion, with the initial expansion round TG-based and the final two rounds PAAG-based (**Supplementary Fig. 5**).

## DISCUSSION

We have found that tetra-gels (TG) made from non-free-radically assembled tetrahedral monomers are capable of mediating expansion microscopy (ExM), and compatible with iterative expansion microscopy (iExM) strategies. When TG-based iExM protocols are compared to classical PAAG-based iExM protocols, using HSV-1 virion envelope proteins as an assay, we see a reduction in the spatial error introduced by the expansion process, from 14.3 nm of added spatial deviation, to 9.2 nm. This raises the tantalizing question of whether more modern hydrogel chemistries such as TG, in an ExM context, might be able to achieve resolutions approaching that of the size of individual biomolecules such as proteins.

Our current study is focused on the chemical principles of the hydrogels of ExM, and is not yet a protocol for general scientific use. For TG-based ExM to be useful in everyday scientific investigations, a number of future improvements will be required. First of all, the chemicals must be made broadly available, through commercialization or other arrangements. We observed TG-based iExM expansion factors ranging from 10x-16x in the current paper, suggesting that refinement of the chemicals to the point of being useful in everyday biology may be of use. Second, in the current TG-based expansion protocol, biomolecules of interest are anchored to the ends of monomers incorporated into the hydrogel network by azide-based click chemistry, meaning that every anchored biomolecule results in a “defect” in the polymer network because it terminates one of the monomer’s arms; replacing such terminal-based anchoring with sidechain-based anchoring via, for example, carbodiimide (EDC) conjugation of proteins to the polyacrylate arms, may improve the final structural homogeneity of the TG polymer network for general ExM purposes. Third, the ideal size of the monomer is not yet established; the monomers are much larger than the sodium acrylate monomers that are used in conventional ExM protocols, raising the question of how well TG monomers permeate dense cells and tissues, and whether smaller TG monomers may be better for expansion of cells and tissues (in contrast to the virions here used for spatial error validation, where the monomers must only bind to the outer motifs of the virions). Finally, appreciation of the fine-scale improvements offered by TG over PAAG most likely will require post-expansion antibody staining, or the use of very small tags such as nanobodies for pre-expansion staining, since the improvements are smaller than the sizes of antibodies; here, we overcame this limit by directly labeling proteins in a nonspecific fashion, but most investigators will want to label specific proteins with well-defined antibodies. To enable post-expansion antibody staining, we will have to replace the disulfide in the monomer with another cleavable moiety that is compatible with the high-temperature (70-121°C) treatments, basic (9-11) pHs, and other harsh conditions that are typically used to denature proteins and expand them away from each other, thus enabling post-expansion antibody staining of endogenous biomolecules^4,8,11^. Alternatively, intramolecular epoxide cross-linking might be helpful for such epitope preservation^43^. Such an expansion process, in the future, may be useful for investigating the detailed nanoscale spatial arrangement of multiple molecular species in clusters and complexes, in cells and tissues, in healthy and disease states.

## METHODS

Methods described here are a concise summary. A more detailed description of each section can be found in the **Supplementary Methods**.

### Synthesis of tetra-gel (TG) monomers

Monomer **1** was synthesized using a procedure modified from a previously described synthesis (**Supplementary Fig. 1**)^27^. First, tetra-arm poly(t-butyl acrylate) with bromo terminal groups (**4**) was synthesized by atom transfer radical polymerization (ATRP). Next, tetra-arm poly(t-butyl acrylate) with azide terminal groups (**5**) was synthesized by replacing bromines of **4** with azides. Lastly, monomer **1** was synthesized by hydrolysis and neutralization of **5** to a final pH of ~7. Monomers **2’**, **2’’**, and **2’’’** were synthesized by N-hydroxysuccinimide (NHS) ester conjugation of the alkynes (DBCO-NHS, BCN-NHS, or DBCO-SS-NHS) to the terminal primary amines of tetra-arm PEGs.

### Cell culture

HEK293FT cells were cultured in chambered coverglasses to a confluency of 60-80%^1^, fixed, and immunostained for expansion microscopy^1,4,8^. Briefly, the cells were treated with 3% (w/v) formaldehyde and 0.1% (w/v) glutaraldehyde in phosphate buffered saline (PBS) for 10 min at room temperature before the subsequent quenching, blocking, and immunostaining steps. HeLa cells were plated on coverglasses coated with Matrigel to a confluency of 50-90% and fixed^1,4,8,14^. Briefly, the cells were treated with 1x PBS + 3% (w/v) formaldehyde + 0.1% (w/v) glutaraldehyde for 10 min at room temperature before the subsequent quenching and blocking steps.

### Thy1-YFP mouse brain slice

All procedures involving Thy1-YFP mice were in accordance with the US National Institutes of Health Guide for the Care and Use of Laboratory Animals and approved by the MIT Committee on Animal Care. 50-100 μm coronal brain slices of Thy1-YFP transgenic mice of 2-4 months old, both male and female, were prepared and immunostained for expansion^4,8,13^.

### General procedure for gelation, digestion, and expansion

Fixed (and immunostained) cells and tissues were incubated in ~0.1-0.2 mg/mL NHS-azide in PBS overnight at room temperature and washed with PBS twice. For the gelling solution, the two monomer solutions were mixed at a targeted molar ratio of 1:1, and an additional amount of water was added to adjust the final concentration of monomer **1** to ~3.3 % (w/v). For example, 10 µL each of monomer **1** and monomer **2’** (both ~200 mg/mL) and 40 µl of water were mixed to yield a gelling solution. Gelation was carried out for 1-2 hours at 37 °C (blank gels) or overnight at 4 °C (cell and tissue samples) in a gelation chamber^4,8^. The gelled cell and tissue samples were incubated in the digestion buffer with proteinase K (8 units/mL) overnight at room temperature^4,8^ and expanded in an excess amount of water three times, each time for 20 min.

### Expansion of HeLa cells (pre-expansion immunostaining and iterative expansion)

Briefly, fixed HeLa cells were stained with primary antibodies, oligo-conjugated secondary antibodies, and azide-modified tertiary oligos as previously described^1,14^. The cells were gelled with a cleavable TG gelling solution prepared by mixing monomer **1**, monomer **2’’’**, and water. The gelled samples were incubated in digestion buffer with Proteinase K at 8 U/mL overnight at room temperature with gentle shaking before de-hybridization of the oligos from the gel-anchored oligos. The expanded samples were re-embedded in N,N′-bis(acryloyl)cystamine (BAC)-crosslinked non-expanding PAAG, hybridized with 1^st^ linker oligos, re-embedded in N,N′-diallyl L-tartardiamide (DATD)-crosslinked expanding PAAG, and incubated in BAC-cleaving buffer. For fluorescence readout, the samples were incubated with fluorophore-conjugated LNA oligos and expanded in water.

### Expansion of HSV-1 virions (direct-labeling and iterative expansion)

Purified HSV-1 virion stock^44^ was diluted before being drop-casted onto a plasma cleaned #0 circular 12-mm coverslip. After 15 min of incubation at room temperature, the virions were fixed in 4% PFA in PBS for 10 min. Azide-modified oligos were directly conjugated to the virion envelope proteins via SoluLink bioconjugation as previously described^1^. The virions were gelled, digested, expanded, and hybridized with fluorophore-conjugated LNA oligos using a similar procedure as for the HeLa cell expansion. For 3-round iterative expansion, BAC-cleaved samples were re-embedded in DATD-crosslinked non-expanding PAAG, hybridized with 2^nd^ linker oligos, re-embedded in bis-crosslinked expanding PAAG, and incubated in DATD-cleaving buffer before the LNA oligo-conjugation, expansion, and imaging. For the PAAG-control, see Supplementary Methods.

### Imaging

All the expanded samples were imaged with a diffraction-limited spinning disk confocal microscope with a 40x, 1.15 NA water-immersion objective. The two-color HeLa cell images and the HSV-1 virion images were deconvolved with theoretical point-spread-functions (PSFs) before visualization and image analysis.

### Image analysis

Averaged HSV-1 virion particles were generated using a semi-automated image analysis pipeline (“Particle Analysis Assistant”)^45^. Each virion particle was manually aligned, automatically cropped, calibrated with the expansion factor, and arithmetically averaged. Using Particle Analysis Assistant, roundness (deviation from the perfect circle) of the HSV-1 virion particle was measured using the standard deviation of the radii within each virion particle. First, the radii in 8 directions (45 degrees apart) were measured for each virion particle as the distance from the particle centroid to the Gaussian-fitted center of the envelope profile. After inspection to remove unfitted profiles, the standard deviation of the accepted radii within the same particle was calculated. For normalization, accepted radii from the same virion particle were normalized by their mean.

## Acknowledgements

We thank Y.-Y. Chou and T. Kirchhausen at HMS for help with VSV stock preparation and virion immobilization; C. Linghu, and O. Shemesh for help with HSV-1 stock preparation; P. Valdes and C. Zhang for helpful discussion about sample staining and expansion; M.J. Kauke for helpful discussion about DNA staining; S.M. Asano for helpful discussion about image analysis; G.H. Huynh for help with mouse brain slice preparation; F. Chen for helpful discussion about monomer and polymerization chemistry design; R. Herlo at HMS for helpful discussion about virion envelope proteins. E.S.B. acknowledges Lisa Yang and Y. Eva Tan, John Doerr, the Open Philanthropy Project, NIH 1R01NS087950, NIH 1RM1HG008525, NIH 1R01DA045549, NIH 2R01DA029639, NIH 1R01NS102727, NIH 1R01EB024261, NIH 1R01MH110932, the HHMI-Simons Faculty Scholars Program, IARPA D16PC00008, U.S. Army Research Laboratory and the U.S. Army Research Office under contract/grant W911NF1510548, U.S.-Israel Binational Science Foundation Grant 2014509, NSF Grant 1734870, and NIH Director’s Pioneer Award 1DP1NS087724. C.-C.Y. acknowledges the McGovern Institute for Brain Research at MIT for the Friends of the McGovern Fellowship. S.U. gratefully acknowledges the Fellows program of the Image and Data Analysis Core at Harvard Medical School.

## Author contributions

R.G. and L.G. designed and synthesized the monomers and conducted initial gelation experiments. C.-C.Y. and R.G. designed and conducted iterative expansion, virion expansion, and associated analyses. C.-C.Y. created the semi-automated virion analysis pipelines. K.D.P. helped characterization of the gel in cell culture. R.L.N. purified HSV-1 and prepared the virion stock solution. S.U. provided purified VSV and conducted initial virion immobilization experiments. C.-C.Y., R.G., and L.G. processed and performed quantitative analysis of all image data. R.G., C.-C.Y., and E.S.B. wrote the manuscript with input from all co-authors. E.S.B supervised the project.

