## Supplementary Material for "A highly homogeneous expansion microscopy polymer composed of tetrahedron-like monomers"

**Supplementary Information**

Supplementary Methods
Supplementary Figs. 1-5
Supplementary Tables 1-2

### SUPPLEMENTARY METHODS

#### Synthesis of tetra-arm sodium poly-acrylate with azide terminal groups (1) (monomer 1)

Monomer **1** was synthesized using a modified procedure as previously described[27] (**Supplementary Fig. 1**). Unless otherwise noted, all chemicals were purchased from Sigma Aldrich.

First, tetra-arm poly(t-butyl acrylate) with bromo terminal groups (**4**) was synthesized by atom transfer radical polymerization (ATRP). Before the synthesis, t-butyl acrylate was purified with an inhibitor removal column to remove a trace amount of 4-methoxyphenol. Next, 128 mL of the purified t-butyl acrylate was added to 640 mg of copper (I) bromide and 48 mg of copper (II) bromide, and the mixture was bubbled with dry nitrogen at 50 °C before 1.03 mL of N,N,N′,N″,N″-pentamethyl diethylenetriamine (PMDETA) was added dropwise. After 5-10 min of continuous bubbling and stirring, a 16 mL acetone solution of 1.6 g pentaerythritol tetrakis(2-bromoisobutyrate) (**3**) was added dropwise. The reaction mixture was stirred at 50 °C for 90 min, with dry nitrogen bubbling on for the first ~10 min. After the reaction, unreacted t-butyl acrylate was removed by rotary evaporation before the crude product mixture was dissolved in dimethylformamide (DMF) and precipitated with water. The precipitation was repeated two to three times additionally, yielding 15.3 g of tetra-arm poly(t-butyl acrylate) with bromo terminal groups (**4**) as a white powder.

Next, tetra-arm poly(t-butyl acrylate) with azide terminal groups (**5**) was synthesized by replacing bromines of **4** with azides. 15.3 g of **4** was dissolved in 80 mL of DMF and an excess amount of sodium azide (exceeding its solubility in DMF) was added to the reaction mixture. The reaction was carried out overnight at room temperature and the supernatant was subsequently decanted and precipitated with water, yielding 11.0 g of tetra-arm poly(t-butyl acrylate) with azide terminal groups (**5**) as a white powder.

Finally, monomer **1** was synthesized by hydrolysis and neutralization of **5**. A total of 5.04 g of **5** was dissolved in 30 mL of methylene chloride, followed by addition of 15 mL of trifluoroacetic acid (TFA). The hydrolysis reaction was carried out at 4 °C with gradual precipitation of a white powder. After 24-48 hours, the precipitated product was collected by centrifugation, washed with acetone, and dried in a low-humidity chamber. The product was re-suspended in an aqueous solution of sodium hydroxide to yield monomer **1** solution (~200 mg/mL) at a final pH of ~7.

#### Synthesis of tetra-arm polyethylene glycol (PEG) with dibenzocyclooctyne (DBCO, 2’), bicyclo[6.1.0]non-2-yne (BCN, 2’’), and dibenzocyclooctyne-disulfide (DBCO-SS, 2’’’) terminal groups (monomer 2)

DBCO-, BCN- and DBCO-SS-terminated monomers **2’**, **2’’**, and **2’’’** were synthesized by N-hydroxysuccinimide (NHS) ester conjugation of the alkynes to the terminal primary amines of tetra-arm PEGs. First, amine-terminated tetra-arm PEG (~10 kDa for **2’** and **2’’**; ~20 kDa for **2’’’**; NOF Corp.) was dissolved in dimethyl sulfoxide (DMSO) to a concentration of 100-200 mg/mL. DBCO-NHS, BCN-NHS or DBCO-SS-NHS was then added to the DMSO solution of the tetra-arm PEG at a 1:1 molar ratio to the total number of terminal amines. Finally, the conjugation reaction was carried out overnight at room temperature to the yield monomer **2'**, **2''**, or **2'''** solutions.

**HEK293 cells**

HEK293FT (Thermo Fisher) cells were cultured in chambered coverglasses (CultureWell, Thermo Fisher) to a confluency of 60-80% as previously described[1]. The cells were then fixed and immunostained[1,4,8]. Briefly, the cells were treated with 3% (w/v) formaldehyde and 0.1% (w/v) glutaraldehyde in phosphate buffered saline (PBS) for 10 min, quenched with 0.1% $NaBH_4$ (w/v) in PBS for 7 min, and then with 100 mM glycine in PBS for 10 min. Immediately after fixation, the cells were permeabilized with 0.1% (w/v) Triton X-100 in PBS for 15 min and blocked with a blocking buffer [5% (v/v) normal donkey serum (NDS) and 0.1% (w/v) Triton X-100 in PBS] for 15 min. For primary antibody staining, the cells were incubated in rat anti-α-tubulin antibody (MA1-80017, Thermo Fisher) solution (1:200 dilution with the blocking buffer) overnight and were washed with the blocking buffer three times, each time for 5 min. For secondary antibody staining, the cells were incubated in Alexa Fluor 488-conjugated donkey anti-rat antibody (A-21208, Thermo Fisher) solution (1: 200 dilution with the blocking buffer) for 5 hours and were washed with the blocking buffer three times, each time for 5 min. Finally, the cells were washed with PBS once for 5 min and stored in PBS for subsequent expansion processes. Unless otherwise noted, all the incubation and washing steps were carried out at room temperature.

**HeLa cells**

HeLa cells were plated on coverglasses coated with Matrigel (BD Sciences) to a confluency of 50-90% and then fixed[1,4,8,14]. Briefly, cells were treated with 1x PBS + 3% (w/v) formaldehyde + 0.1% (w/v) glutaraldehyde for 10 min, quenched with 1x PBS + 0.1% (w/v) $NaBH_4$ for 7 min. The cells were then washed once with 1x PBS + 100 mM glycine for 5 min, and then twice with 1x PBS for 5 min each. Fixed cells were stored at 4°C until the immunostaining step. Unless otherwise noted, all the incubation and washing steps were carried out at room temperature.

**Thy-1 YFP mouse brain slices**

Coronal brain slices of transgenic mice expressing cytosolic YFP under the Thy1 promoter (Thy1-YFP-H strain, Jackson Laboratory) were prepared and immunostained[4,8,13]. Unless otherwise noted, all the incubation and washing steps were carried out at room temperature. Briefly, Thy1-YFP mice, 2-4 months old, both male and female, were anesthetized with ketamine/xylazine and perfused transcardially with 25 mL ice cold 4% (w/v) paraformaldehyde (PFA) in PBS, followed by 25 mL of ice cold PBS. The brains were dissected out and soaked in 4% (w/v) PFA in PBS for 24 hours at 4ºC. 50-100 µm coronal slices were prepared using a vibratome (Leica VT1000S) and were stored in PBS at 4°C for subsequent expansion processes.

For pre-expansion staining, the fixed brain slices were permeabilized and blocked with a blocking buffer [5% (v/v) NDS and 0.1% (w/v) Triton X-100 in PBS] for 2 hours. For primary antibody staining, the slices were incubated with rabbit anti-Homer1 antibody (160003, Synaptic Systems) solution (1: 200 dilution with the blocking buffer) overnight and washed with the blocking buffer twice, each time for 30 min. For secondary antibody staining, the slices were incubated with dye-conjugated Alexa Fluor 647, Cy5, or Alexa Fluor 680 goat anti-rabbit antibody (A21245, A10523, or A21109, Thermo Fisher) solution (1: 200 dilution with the blocking buffer) overnight and washed with the blocking buffer twice, each time for 30 min. Finally, the slices were washed once with PBS and stored in PBS for subsequent expansion processes.

For post-proteolysis staining, the gelled and digested brain slices were incubated with chicken anti-GFP antibody (A10262, Thermo Fisher) solution (1:200 dilution with the blocking buffer) overnight and subsequently with Alexa Fluor 488-conjugated goat anti-chicken antibody (A11039, Thermo Fisher) solution (1:200 dilution with the blocking buffer) overnight before the same washing and storage steps with PBS as described.

**Gelation, digestion, and expansion of cells and tissues (single-round expansion)**

Unless otherwise noted, single-round expansion of cells and tissues with tetra-gels (TGs) were carried out using the following general procedure. First, fixed (and immunostained) cells and tissues were incubated in ~0.1-0.2 mg/mL NHS-azide in PBS overnight at room temperature and washed with PBS twice immediately before gelation.

Next, monomer **1** and monomer **2** solutions were mixed at close to 1:1 molar ratio, and an additional amount of water was added to adjust the final concentration of monomer **1** to ~3.3 % (w/v). In a typical gelation with monomer **2’**, 10 µL of monomer **1** (~200 mg/mL), 10 µL of monomer **2** (~200 mg/mL), and 40 µl of water were mixed to yield the gelling solution. We note that the molar ratio between the mixed monomer **1** to **2’** can vary slightly according to the molecular weight of monomer **1**, which depends on the final conversion rate of the t-butyl ester hydrolysis in the last deprotection step of the synthesis. For example, the exact molar ratio of monomer **1** to **2’** can be ~0.99 at 50% conversion and ~1.16 at 100% conversion. However, due to the complementary, mutually-limiting polymerization mechanism of TG, this slight variation should not significantly alter the composition of the incorporated monomers **1** and **2’**. After drop-casting the gelling solution to the samples in a gelation chamber as previously described[4,8], gelation was carried out for 1-2 hours at 37 °C (blank gels) or overnight at 4 °C (cell and tissue samples). The total amount of monomers being mixed was adjusted proportionally according to the size and number of the samples.

Finally, the gelled cell and tissue samples were incubated in the digestion buffer with proteinase K (8 units/mL) overnight at room temperature as previously described[4,8]. For expansion, the digested samples were washed in an excess amount of water three times, each time for 20 min.

For fluorescein visualization of blank gels, a trace amount of fluorescein amine was mixed into the gelling solution. Briefly, a stock solution of ~50 mM fluorescein-azide was prepared by adding 5 µL of 100 mM fluorescein-amine in DMSO to 5 µL of 20 mg/mL NHS-azide in DMSO. ~3 µL of the fluorescein-azide stock solution was added to ~60 µL of the gelling solution (with monomers **2’**, **2’’**, or **2’’’**). The gelling solution with fluorescein was drop-cast into a circular mold of ~3 mm diameter before gelation for 1-2 hours at 37 °C.

**Expansion of HeLa cells (pre-expansion immunostaining and iterative expansion)**

*Pre-expansion immunostaining*

Fixed cells were stained with primary antibodies, oligo-conjugated secondary antibodies, and azide-modified tertiary oligos (for TG) as previously described[1,14]. Briefly, fixed cells were permeabilized and blocked with HeLa staining buffer [1x PBS + 5% (v/v) normal donkey serum + 0.2% (w/w) Triton X-100] for 10 min. Primary antibody staining was performed with HeLa staining buffer for 1 hr at RT, followed by 3 washes with 1x PBS for 5 min each at RT. Secondary antibody staining was performed with hybridization buffer [10% w/v dextran sulfate, 1 mg/mL yeast tRNA, 5% v/v normal donkey serum, 2x SSC, 0.1% (w/w) Triton X-100] for 1 hr at RT, followed by 3 washes with 1x PBS for 5 min each at RT. Tertiary oligo hybridization was performed by first incubating the sample in hybridization buffer for 3 hr, followed by incubation with the tertiary oligo overnight at RT. After hybridization, samples were washed with 1x PBS 3 times for 5 min each at RT. For single-color tubulin staining, the cells were stained with rabbit anti-beta-tubulin (ab6046, Abcam, 1:100 dilution) primary antibody, and then with oligo-conjugated anti-rabbit secondary antibody (oligo sequence B1'; see Supplementary Table 1 for oligo sequences used), followed by hybridization with azide-modified complementary oligo (oligo sequence B1, at 100 nM). For dual-color tubulin and clathrin staining, the cells were stained with sheep anti-alpha/beta tubulin (ATN02, Cytoskeleton, Inc, 1:50 dilution) and rabbit anti-clathrin heavy chain (ab21679, Abcam, 1:500 dilution) primary antibodies, and then with oligo-conjugated anti-sheep (oligo sequence A2') and anti-rabbit (oligo sequence B1') secondary antibodies, followed by hybridization with azide-modified complementary oligos (oligo sequences B1 and A2, at 100 nM each). Oligo conjugation to secondary antibodies were performed with the same protocol as previously described[1] (also available on www.expansionmicroscopy.org), with the modification that the S-HyNic reaction was performed with 3 times of its concentration (600 µM instead of 200 µM) compared to the original protocol.

*Gelation and digestion*

As described in the previous section, the cleavable TG gelling solution was prepared by mixing monomer **1** (200 mg/mL) and monomer **2'''** (200 mg/mL) at a molar ratio of close to 1:1 and then adding water to adjust the final concentration of monomer **1** to ~3.3% (w/v). A gelation chamber was constructed around the cell-immobilized coverslip using the following steps. First, the coverslip was transferred to the center of a glass slide. Spacers consisting of a stack of a #0 and a #1 coverslip were placed on either side of the cell-immobilized coverslip. 50 µL of freshly prepared TG gelling solution was added to the coverslip, and the chamber was closed by placing a rectangular coverslip on top of the spacers. The gelling solution was further added from the side of the chamber until the chamber was completely filled. Gelation chambers were then placed in a humidified chamber and incubated at 4°C overnight. After the incubation, the chamber was partially opened using a diamond scribe to remove portions of the top cover glass that were not directly above the cell-immobilized coverslip. The chamber was then placed into a rectangular 4-well dish and incubated in digestion buffer with Proteinase K at 8 U/mL (New England Bio Labs; 1:100 dilution) overnight at room temperature with gentle shaking. The diameter of the gel was measured for downstream estimation of the overall expansion factor. Regions inside the circular 12-mm coverslip (i.e. regions with the immobilized cells) were trimmed into a parallelogram, of which the side lengths were measured for downstream estimation of the overall expansion factor. Finally, trimmed gels were washed twice in PBS, each time for 10 min. To de-hybridize B1' and A2' oligos (conjugated to the secondary antibodies) from the gel-anchored B1 and A2 oligos (the azide-modified tertiary oligos), the gels were incubated in 80% formamide at room temperature for 1 hr with gentle shaking, and then washed three times in PBST (1x PBS + 0.1% Triton X-100) at room temperature with gentle shaking, for 30 min each.

*Re-embedding into a BAC-crosslinked non-expanding 2nd gel*
The gels were transferred (with the cell side facing down) into a rectangular 4-well dish that carries a glass slide in each well, and expanded in water three times, each time for 30 min. The gels were then incubated in 3 mL of BAC-crosslinked non-expanding gelling solution [10.4% (w/v) acrylamide, 0.2% (w/v) BAC, 0.05% (w/v) TEMED, 0.05% (w/v) APS] for 5 min with gentle shaking. After the incubation, the non-expanding gelling solution was removed from the well, and the glass slide carrying the expanded gel was transferred to a gelation chamber. Spacers consisting of a stack of two #1.5 cover glasses were placed on either side of the gel, and the chamber was closed with a rectangular cover glass. The non-expanding gelling solution was added from the side of the chamber until the chamber was completely filled. The gelation chambers were incubated for 2 hours at 37°C. After gelation, the chambers were opened by removing the top cover glass. Side lengths of the parallelogram were measured. The gels were trimmed to leave only the portion inside the parallelogram, while preserving the shape of the parallelogram. Side lengths of the trimmed gels were measured. Finally, the trimmed gels were washed twice in PBS, each time for 30 min.

*1st Linker hybridization*
The gels were incubated in hybridization buffer (4x SSC + 20% (v/v) formamide) for 30 min at room temperature. The gels were incubated with 1 nmol of oligo 5'Ac-B1'-4xB2' in 1 mL of hybridization buffer overnight at room temperature. After incubation, the gels were washed in hybridization buffer three times, each time for 1 hour, and then overnight, all with gentle shaking. The gels were then washed three times in PBS, each time for 5 min.

*Re-embedding into a DATD-crosslinked expanding 3rd gel*
The gels were incubated in DATD-crosslinked expanding gelling solution [8.6% (w/v) sodium acrylate, 2.6% (w/v) acrylamide, 0.5% (w/v) DATD, 1x PBS, 2M NaCl, 0.01% (w/v) 4-HT, 0.2% (w/v) TEMED, 0.2% (w/v) APS] for 30 min at 4°C. The gels (with the cell side facing down) were then enclosed in gelation chambers, incubated for 2 hours at 37°C, size-measured, trimmed, size-re-measured, and washed as described in "*Re-embedding into a BAC-crosslinked non-expanding 2nd gel*".

*Cleaving BAC-crosslinked 1st and 2nd gels*
The gels were incubated in BAC-cleaving buffer (0.25M TCEP-HCl, 0.75M Tris-HCl, pH 8.0) overnight at room temperature. The gels were then washed four times in PBS, each time for 30 min.

*LNA hybridization for readout after 2-round expansion*
The gels were incubated with 1 nmol of fluorophore-conjugated LNA oligo (LNA_B2-Atto647N and/or LNA_A1-Atto565; see sequences in Supplementary Table 1) in 500 µL of hybridization buffer. The LNA-hybridized gels were washed in hybridization buffer three times, each time for 1 hour, and then overnight, all with gentle shaking. The gels were then washed three times in PBS, each time for 5 min.

*Gel expansion, immobilization, and imaging for 2-round expanded samples*
The gels were trimmed into smaller pieces (~5 by 5 mm) while preserving the shape of the parallelogram. First, the gels were transferred (with the cell side facing down) into a rectangular 4-well dish that carries a glass slide in each well, and expanded in water three times, each time for 30 min. A glass-bottom 6-well plate was modified with poly-lysine, and the expanded gels were gently transferred to the poly-lysine modified coverslip surface for imaging, as previously described[4].

**Expansion of HSV-1 virions (direct labeling and iterative expansion)**

*Immobilization and fixation*
Purified HSV-1 virion stock was prepared by the Viral Core Facility at the Massachusetts General Hospital (MGH) as previously described[44]. The HSV-1 stock was diluted to a functional titer of $2.5 \times 10^8$ functional virions/mL in PBS and kept on ice until immobilization. A #0 circular 12-mm coverslip was cleaned with a plasma cleaner (PDC-001, Harrick Plasma) for 1 min. Immediately after the plasma cleaning, 30 µL of the diluted HSV-1 solution was drop-casted onto the coverslip and incubated for 15 min at room temperature. The immobilized virions were fixed in 4% PFA in PBS for 10 min, and then washed with PBS twice, each time for 5 min.

*Oligo conjugation to envelope proteins*
Envelope proteins on the fixed virions were conjugated to DNA oligos with the SoluLink bioconjugation chemistry as previously described[1]. The oligo provided a molecular handle for label anchoring, transfer, and amplification through the iterative expansion process. Briefly, a 22-bp oligo [sequence B1' with a 5' amine modification[14] (Integrated DNA Technologies)] was purified with ethanol precipitation and reacted with Sulfo-S-4FB (S4FB) overnight in Buffer A (150 mM NaCl, 100 mM $Na_2HPO_4$, pH 7.4) at a molar ratio of 1:15. The S4FB-reacted oligo was purified with a size exclusion filter, and then stored at 4°C. Fixed virions immobilized on the coverslip were washed in Buffer A for 5 min, and then incubated with 100 µL of 160 mM S-HyNic in Buffer A for 2 hours at room temperature. The S-HyNic-reacted virions were washed with Buffer C (150 mM NaCl, 100 mM $Na_2HPO_4$, pH 6.0) twice, each time for 5 min. Oligo conjugation solution was prepared by first adding 50 nmol of purified S4FB-reacted oligo to 100 µL of Buffer C, and then adding an amount of 10x TurboLink Catalyst Buffer that equals to 1/9 of the combined volume. The S-HyNic-reacted virions were incubated in the oligo conjugation buffer overnight at room temperature in a humidified chamber. Next, the oligo-conjugated virions were washed three times with PBS, each time for 10 min, and then incubated in detergent-free hybridization buffer (10% (w/v) dextran sulfate, 1 mg/mL yeast tRNA, 5% (v/v) NDS, 2x SSC) for 3 hours at room temperature. The virions were incubated with 4 nmol of oligo B1-acrydite or oligo B1-azide for the subsequent gelation with TGs [or sodium polyacrylate/acrylamide gels (PAAGs)], in 300 µL of the detergent-free hybridization buffer, overnight at room temperature. Finally, the virions were washed three times in PBS, each time for 10 min.

*Gelation and digestion*
As described in the previous section, cleavable TG gelling solution was prepared by mixing monomer **1** (200 mg/mL) and monomer **2'''** (200 mg/mL) at a molar ratio of close to 1:1 and then adding water to adjust the final concentration of monomer **1** to ~3.3% (w/v). BAC-crosslinked cleavable PAAG gelling solution was prepared as previously described[14]. A gelation

chamber was constructed around the virus-immobilized coverslip using the following steps. First, the coverslip was transferred to the center of a glass slide. Spacers consisting of a stack of a #0 and a #1 coverslip were placed on either side of the virus-immobilized coverslip. 50 µL of freshly prepared TG or PAAG gelling solution was added to the coverslip, and the chamber was closed by placing a rectangular coverslip on top of the spacers. The gelling solution was further added from the side of the chamber until the chamber was completely filled. After gelation for 2 hours at 37°C, the chamber was partially opened using a diamond scribe to remove portions of the top cover glass that were not directly above the virus-immobilized coverslip. The chamber was then placed into a rectangular 4-well dish and incubated in digestion buffer with Proteinase K at 8 U/mL (New England Bio Labs; 1:100 dilution) overnight at room temperature with gentle shaking. After digestion, the top cover glass came off naturally and was removed from the solution. The diameter of the gel was measured for downstream estimation of the overall expansion factor. Regions inside of the circular 12-mm coverslip (i.e. regions with the immobilized viruses) were trimmed into a parallelogram, of which the side lengths were measured for downstream estimation of the overall expansion factor. Finally, trimmed gels were washed twice in PBS, each time for 10 min. To de-hybridize B1' and B1 oligos, the gels were incubated in 80% formamide at room temperature with gentle shaking for 1 hr, and then washed three times in PBS, each time for 10 min.

*Re-embedding into a BAC-crosslinked non-expanding 2$^{nd}$ gel*

The gels were transferred (with the virion side facing down) into a rectangular 4-well dish that carries a glass slide in each well, and expanded in water three times, each time for 30 min. The gels were then incubated in 3 mL of BAC-crosslinked non-expanding gelling solution [10.4% (w/v) acrylamide, 0.2% (w/v) BAC, 0.05% (w/v) TEMED, 0.05% (w/v) APS] for 5 min with gentle shaking. After the incubation, the non-expanding gelling solution was removed from the well, and the glass slide carrying the expanded gel was transferred to a gelation chamber. Spaces consisting of a stack of #1.5 cover glasses were placed on either side of the gel, and the chamber was closed with a rectangular cover glass. The non-expanding gelling solution was added from the side of the chamber until the chamber was completely filled. The gelation chambers were incubated for 2 hours at 37°C. After gelation, the chambers were opened by removing the top cover glass. Side lengths of the parallelogram were measured. The gels were trimmed to leave only the portion inside the parallelogram, while preserving the shape of the parallelogram. Side lengths of the trimmed gels were measured. Finally, the trimmed gels were washed twice in PBS, each time for 10 min.

*1$^{st}$ Linker hybridization*

The gels were incubated in hybridization buffer (4x SSC + 20% (v/v) formamide) for 30 min at room temperature. For readout after 2-round expansion (~10-20x expansion factor), the gels were incubated with 1 nmol of oligo 5'Ac-B1'-4xB2' in 500 µL of hybridization buffer overnight at room temperature. For readout after 3-round expansion (~40-80x expansion factor), the gels were incubated with 1 nmol of oligo 5'Ac-B1'-A2' in 500 µL of hybridization buffer overnight at room temperature. After incubation, the gels were washed in hybridization buffer three times, each time for 1 hour, and then overnight, all with gentle shaking. The gels were then washed three times in PBS, each time for 5 min.

*Re-embedding into a DATD-crosslinked expanding 3$^{rd}$ gel*

The gels were incubated in DATD-crosslinked expanding gelling solution [8.6% (w/v) sodium acrylate, 2.6% (w/v) acrylamide, 0.5% (w/v) DATD, PBS, 2M NaCl, 0.01% (w/v) 4-HT, 0.2% (w/v) TEMED, 0.2% (w/v) APS] for 30 min at 4°C. The gels (with the virion side down) were enclosed in gelation chambers, incubated for 2 hours at 37°C, size-measured, trimmed, size-re-measured, and washed as described in "*Re-embedding into a BAC-crosslinked non-expanding 2$^{nd}$ gel*".

*Cleaving BAC-crosslinked 1$^{st}$ and 2$^{nd}$ gels*
The gels were incubated in BAC-cleaving buffer (0.25M TCEP-HCl, 0.75M Tris-HCl, pH 8.0) overnight at room temperature. The gels were then washed four times in PBS, each time for 30 min. For samples designated for 3-round expansion, the gels were incubated in thiol-blocking buffer (100 mM maleimide, 100 mM MOPS, pH 7.0) for 2 hours at room temperature. The thiol-blocked gels were washed three times in PBS, each time for 10 min.

*LNA hybridization for readout after 2-round expansion*
For samples designated for 2-round expansion, the gels were incubated with 1 nmol of LNA_B2-Atto647N in 500 µL of hybridization buffer. The LNA-hybridized gels were washed in hybridization buffer three times, each time for 1 hour, and then overnight, all with gentle shaking.  The gels were then washed three times in PBS, each time for 5 min.

*Gel expansion, immobilization, and imaging for 2-round expanded samples*
The gels were trimmed into smaller pieces (~5 by 5 mm) while preserving the shape of the parallelogram. First, the gels were transferred (with the virion side down) into a rectangular 4-well dish that carries a glass slide in each well, and expanded in water three times, each time for 30 min. A glass-bottom 6-well plate was modified with poly-lysine, and the expanded gels were gently transferred to the poly-lysine modified coverslip surface for imaging, as previously described[4].

*Re-embedding into a DATD-crosslinked non-expanding 4$^{th}$ gel (for 3-round expansion)*
Thiol-blocked gels in "*Cleaving BAC-crosslinked 1$^{st}$ and 2$^{nd}$ gels*" were subsequently trimmed into smaller pieces (~5 by 5 mm) while preserving the shape of the parallelogram. The gels were transferred (with the virion side down) into a rectangular 4-well dish that carries a glass slide in each well, and expanded in water three times, each time for 30 min. The gels were transferred onto a slide glass and trimmed in the z-direction into a thickness of 1 mm. Briefly, the glass slide (with 1-mm thickness) carrying the expanded sample was placed between two stacks of 1-mm-glass slides, and a cryostat blade was pushed slowly through the expanded gel. The bottom gel, which carries the virus at the bottom side, was transferred back to the 4-well plate. The z-trimmed gels were then incubated in DATD non-expanding gelling solution [10.4% (w/v) acrylamide, 0.5% (w/v) DATD, 0.05% (w/v) TEMED, 0.05% (w/v) APS] for 30 min at 4°C. The gels were enclosed in gelation chambers, incubated for 2 hours at 37°C, size-measured, trimmed, size-re-measured, and washed as described in "*Re-embedding into a BAC-crosslinked non-expanding 2$^{nd}$ gel*".

*2$^{nd}$ Linker hybridization*
The gels were incubated in hybridization buffer (4x SSC + 20% formamide) for 30 min at room temperature. The gels were incubated with 0.5 nmol of oligo 5'Ac-A2-4xB2' in 1 mL of

hybridization buffer overnight at room temperature. After incubation, the gels were washed in hybridization buffer three times, each time for 1 hour, and then overnight, all with gentle shaking. The gels were then washed twice in PBS, each time for 30 min.

*Re-embedding into a bis-crosslinked expanding 5th gel*
The gels were incubated in bis-crosslinked expanding gelling solution [8.6% (w/v) sodium acrylate, 2.6% (w/v) acrylamide, 0.15% (w/v) N,N-methylenebisacrylamide (bis), PBS, 2M NaCl, 0.01% (w/v) 4-HT, 0.2% (w/v) TEMED, 0.2% (w/v) APS] for 30 min at 4°C. The gels (with the virus side down) were enclosed in gelation chambers, incubated for 2 hours at 37°C, size-measured, trimmed, size-re-measured, and washed in the same way as described in "*Re-embedding into a BAC-crosslinked non-expanding 2nd gel*".

*Cleaving DATD-crosslinked 4th and 5th gels*
The gels were incubated in DATD-cleaving buffer (20 mM sodium periodate, PBS, pH 5.5) for 30 min at room temperature. The gels were then washed three times in PBS, each time for 30 min, and then overnight with gentle shaking.

*LNA hybridization for readout after 3-round expansion*
The gels were hybridized with LNA_B1_Atto647N as described in "*LNA hybridization for readout after 2-round expansion*".

*Gel expansion, immobilization, and imaging*
The gels were trimmed, expanded, immobilized and imaged as described in "*Gel expansion, immobilization, and imaging for 2-round expanded samples*".

*Expansion factor estimation*
Side lengths of the gels were recorded before and after each trimming step (for example, after every re-embedding step and before every immobilization step) and immediately before imaging. Single-stage expansion factor was calculated by taking the averaged quotient between the pre-trimming size of the current step and the post-trimming size of the previous step. Overall expansion factor was calculated from the product of all the previous single-step expansion factors until the final step.

### Imaging

Unless otherwise noted, all the expanded samples were imaged with an Andor spinning disk (CSU-W1, Yokogawa) confocal system on a Nikon Eclipse Ti-E microscope body with a CFI Apo LambdaS LWD 40x, 1.15 NA water-immersion objective (Nikon). The two-color HeLa cell tubulin and clathrin images and the HSV-1 virion images were deconvolved with the theoretical point-spread functions (PSFs) (Huygens Essential, SVI) before visualization and image analysis.

### Visualization

Unless otherwise noted, all 3D renderings were generated using Imaris x64 8.3 (Oxford Instruments).

### Averaged HSV-1 virion images

An averaged HSV-1 virion particle was generated using a semi-automated image analysis pipeline implemented on MATLAB ("Particle Analysis Assistant"). The "Particle Analysis Assistant" is available for download[45]. First, within an acquired image z-stack, all round objects with a local minimum inside the object were identified as virion particles. The center of each virion was then determined manually within the image z-slice that had the largest virion diameter. Next, the center of the virions was re-inspected and re-aligned once more. During the second inspection, a small portion (<10%) of the virions, which had significant overlaps with the neighboring virions, were rejected from the averaging. Finally, the single virion images around each virion center were automatically cropped, calibrated with the expansion factor, and arithmetically averaged.

#### HSV-1 virion envelope protein layer roundness analysis

Roundness (deviation from the perfect circle) of the HSV-1 virion envelope protein layer was measured by the standard deviation of the radii within each virion particle, using the semi-automated image analysis pipeline implemented on MATLAB ("Particle Analysis Assistant")[45]. All HSV-1 virion particles that passed the second inspection in the previous section were analyzed with the following semi-automated procedure. For each particle, the z-plane that corresponded to the vertical center of the particle was manually identified by selecting the one with the maximum contour diameter of the envelope layer. From the centerline z-plane image, the radii in 8 directions (45 degrees apart) were algorithmically measured by computing the distance from the particle centroid to the Gaussian-fitted center of the envelope profile. Manual inspection was carried out to exclude portions of the 8 radii that were measured either incorrectly or with low confidence, based on the following criteria for rejection: (1) when the line intensity profile contained peaks that belonged to other virion particles, (2) when the envelope layer in the particular direction was not present or was significantly dimmer than in other directions, or (3) when the automated Gaussian fitting failed. Standard deviation of all the accepted radii within the same particle was reported as population statistics. For normalization, the accepted radii were divided by their mean (such that the mean of all the normalized radii was equal to 1).

### SUPPLEMENTARY FIGURES AND TABLES

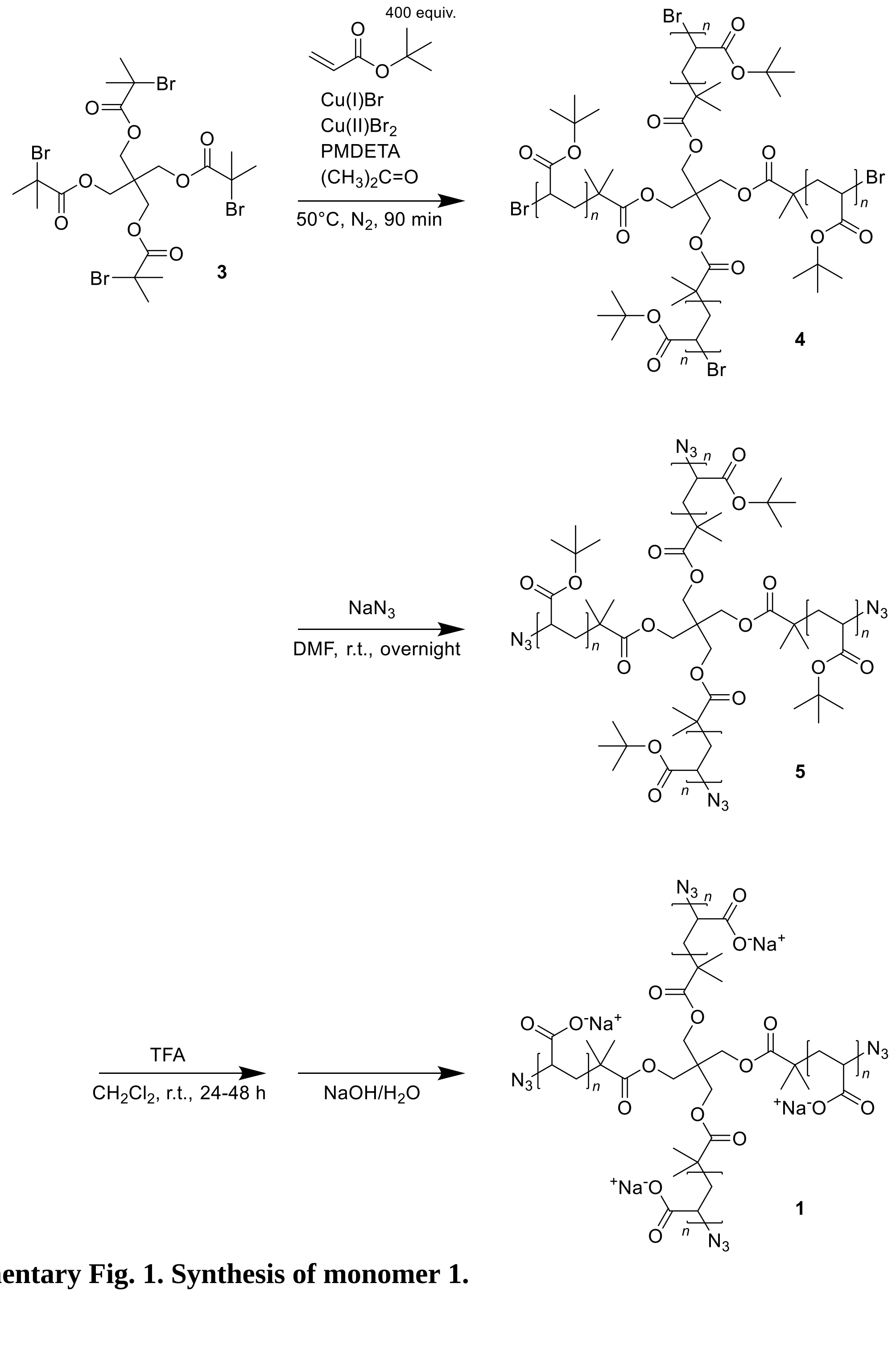


**Supplementary Fig. 1. Synthesis of monomer 1.**

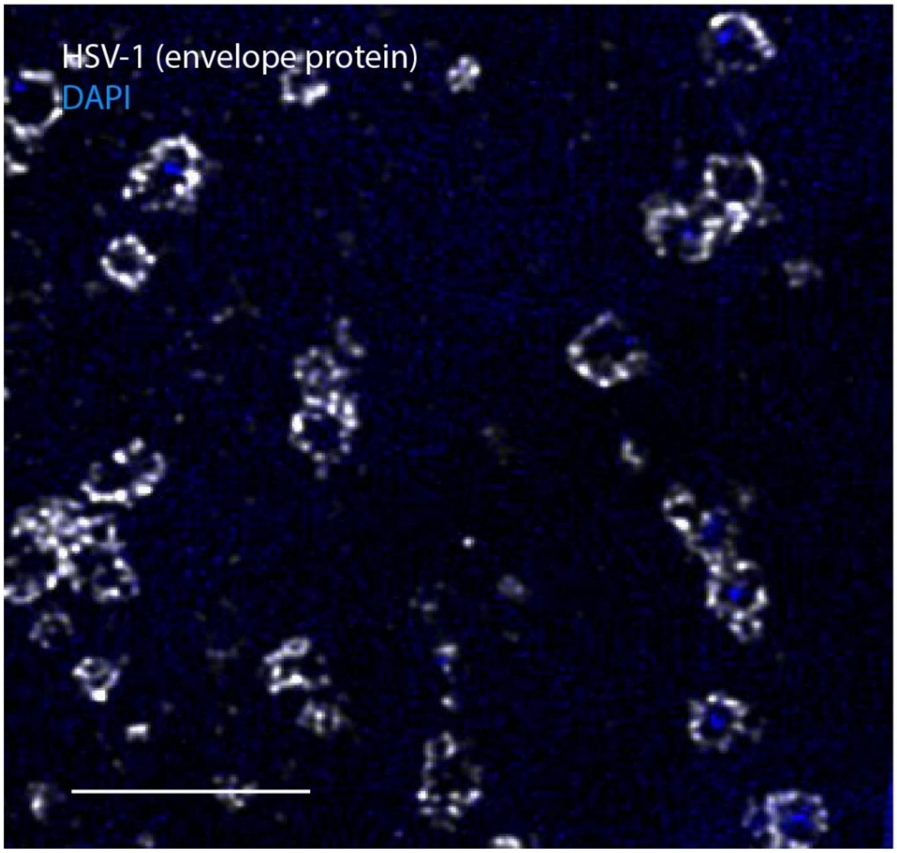


**Supplementary Fig. 2. Expansion and two-color imaging of herpes simplex virus type 1 (HSV-1) envelope proteins (white) and DNA (blue).** The virions were expanded by TG-based 2-round iterative expansion with direct labeling of the envelope proteins. Expansion factor, 10.7x. Scale bar, 1 µm (10.7 µm).

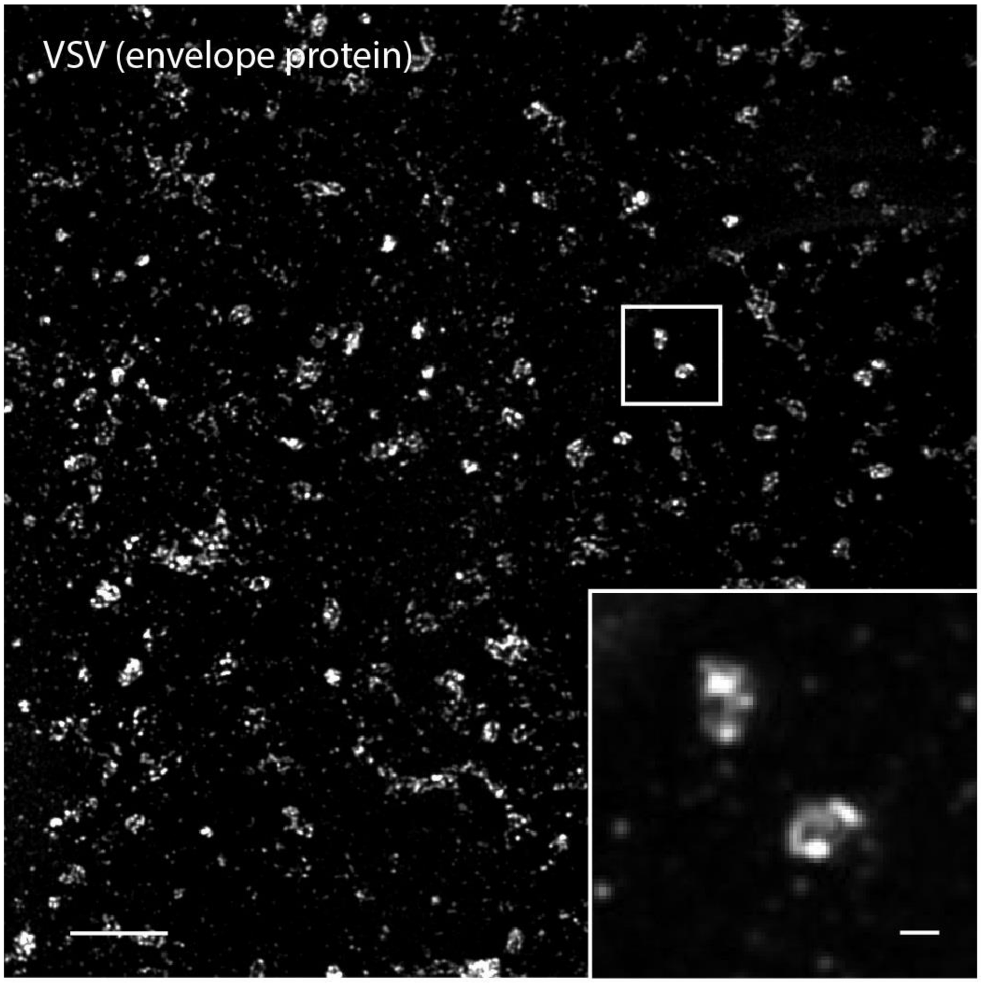


**Supplementary Fig. 3. Expansion of vesicular stomatitis virus (VSV) virions.** The virions were expanded by TG-based 2-round iterative expansion with direct labeling of the envelope proteins. Expansion factor, 10.9x. Scale bar, 1 µm (10.9 µm). Inset, magnified view of the boxed region on a single xy-plane. Scale bar, 100 nm (1.09 µm).

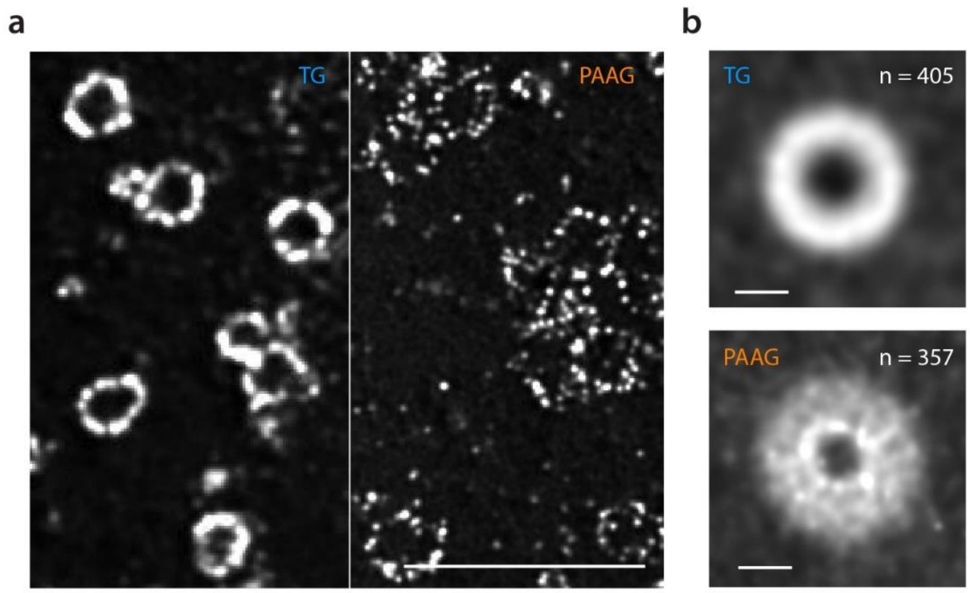


**Supplementary Fig. 4. Expansion of HSV-1 virions. a,** HSV-1 virions with directly labeled envelope proteins, expanded by TG- (left) and sodium polyacrylate/acrylamide gel (PAAG)-based (right) 2-round iterative expansion. Scale bar, 1 µm (TG, 10.3 µm; PAAG, 15.3 µm). Expansion factors, 10.3x (TG) and 15.3x (PAAG). **b,** Averaged single-particle images of HSV-1 virions after TG- (top) and PAAG-based (bottom) 2-round iterative expansion (TG, n = 405; PAAG, n = 357 virion particles; virion particles from the same single batch of live HSV-1 preparation). Scale bars, 100 nm.

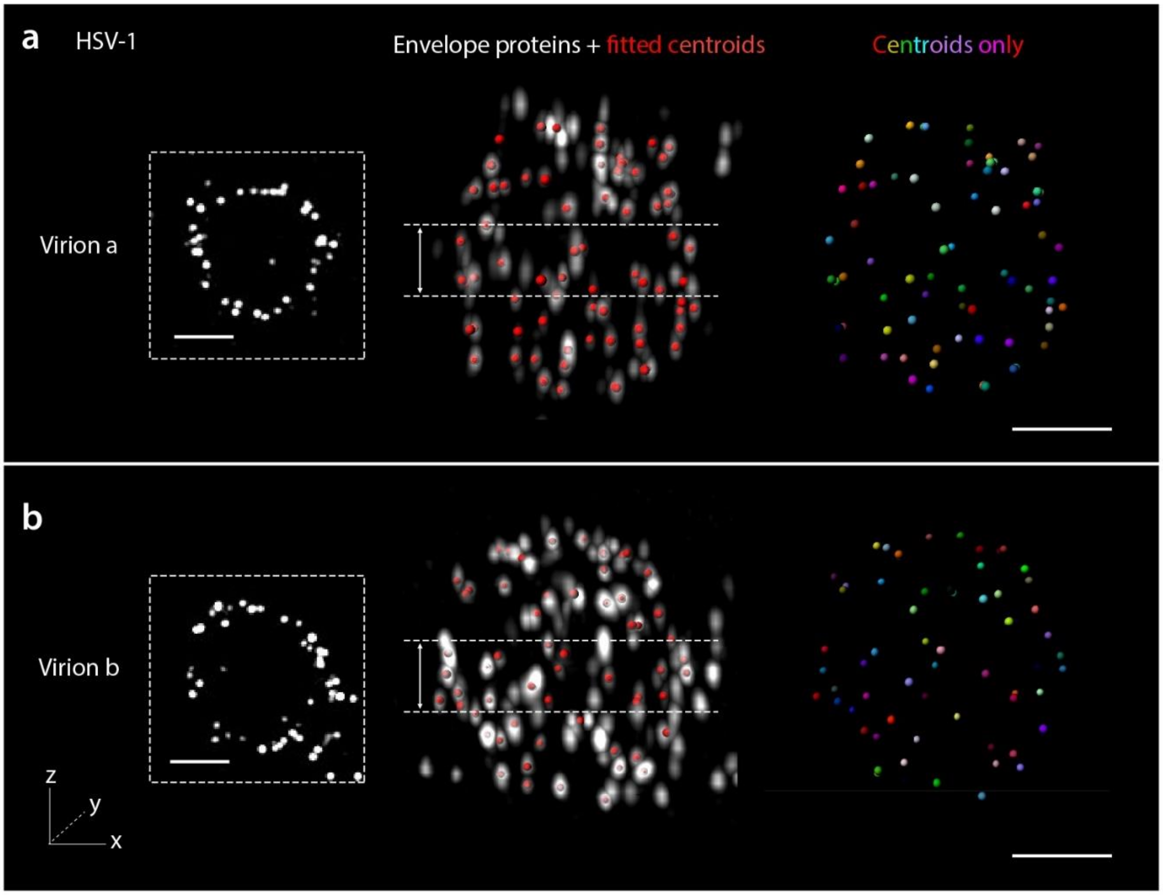


**Supplementary Fig. 5. 3D rendered images of the envelope proteins of two HSV-1 virions expanded by TG-based 3-round iterative expansion with direct labeling.** Overlaid images of the deconvolved puncta (white) and the fitted centroids (red) are shown on the left, and the extracted centroids (colored) are shown on the right. Expansion factors, 38.8x (virion a) and 38.3x (virion b). Scale bars, 100 nm (virion a, 3.88 µm; virion b, 3.83 µm). Insets (dotted boxes), maximum intensity projection (MIP) of the same virions over a ~65 nm range around the particle center (between the dotted lines in the 3D rendered images). Scale bars, 100 nm (virion a, 3.88 µm; virion b, 3.83 µm).

**Supplementary Table 1. DNA oligo sequences.**

| Oligo Name | Purpose | Sequence (IDT format) | Modification |
|---|---|---|---|
| For iterative expansion of HeLa cells (Fig. 3) | | | |
| 5'Amine-B1' | Conjugation to secondary antibody for pre-G1 immunostaining | AAT ACG CCC TAA GAA TCC GAA C | 5' Amino Modifier C6 |
| 5'Amine-A2' | Conjugation to secondary antibody for pre-G1 immunostaining | AAG GTG ACA GGC ATC TCA ATC T | 5' Amino Modifier C6 |
| 5'Azide-B1 | Pre-G1 adaptor for TG-based iExM | GTT CGG ATT CTT AGG GCG TA | 5' Azide |
| 5'Azide-A2 | Pre-G1 adaptor for TG-based iExM | AGA TTG AGA TGC CTG TCA CC | 5' Azide |
| 5'Acrydite-B1'-4xB2' | Post-G2 linker | TAC GCC CTA AGA ATC CGA ACA TGC ATT ACA GCC CTC AAT GCA TTA CAG CCC TCA ATG CAT TAC AGC CCT CAA TGC ATT ACA GCC CTC A | 5' Acrydite |
| 5'Acrydite-A2'-4xA1' | Post-G2 linker | GGT GAC AGG CAT CTC AAT CTA TTA CAA AGC ATC AAC GAT TAC AAA GCA TCA ACG ATT ACA AAG CAT CAA CGA TTA CAA AGC ATC AAC G | 5' Acrydite |
| LNA_B2-Atto647N | Post-G3 readout | TGAGGGCTGTAATGC | 3' Atto 647N, LNAs (underlined) |
| LNA_A1-Atto565 | Post-G3 readout | CGTTGATGCTTTGTA | 3' Atto 565, LNAs (underlined) |
| For iterative expansion of HSV virions (Fig. 4) | | | |
| 5'Amine-B1' | Pre-G1 conjugation to envelope proteins | AAT ACG CCC TAA GAA TCC GAA C | 5' Amino Modifier C6 |
| 5'Acrydite-B1 | Pre-G1 adaptor for PAAG-based iExM | GTT CGG ATT CTT AGG GCG TA | 5' Acrydite |
| 3'Azide-B1 | Pre-G1 adaptor for TG-based iExM of virions | GTT CGG ATT CTT AGG GCG TA | 3' Azide |
| 5'Acrydite-B1'-4xB2' | Post-G2 linker for 2-round iExM | TAC GCC CTA AGA ATC CGA ACA TGC ATT ACA GCC CTC AAT GCA TTA CAG CCC TCA ATG CAT TAC AGC CCT CAA TGC ATT ACA GCC CTC A | 5' Acrydite |

| | | | |
|---|---|---|---|
| 5’Acrydite-B1’-A2’ | Post-G2 linker for 3-round iExM | TAC GCC CTA AGA ATC CGA ACA TGG TGA CAG GCA TCT CAA TCT | 5’ Acrydite |
| 5’Acrydite-A2-4xB2’ | Post-G4 linker for 3-round iExM | AGA TTG AGA TGC CTG TCA CCA TGC ATT ACA GCC CTC AAT GCA TTA CAG CCC TCA ATG CAT TAC AGC CCT CAA TGC ATT ACA GCC CTC A | 5’ Acrydite |
| LNA_B2-Atto647N | Post-G3 or Post-G5 readout | TGAGGGCTGTAATGC | 3’ Atto 647N, LNAs (underlined) |

**Supplementary Table 2. Recipes of hydrogel gelling solutions.**

| Gel Name | Purpose | Recipe (w/v, unless otherwise noted) | | | | |
|---|---|---|---|---|---|---|
| Tetra-gel (TG) | | | | | | |
| | | Monomer **2’**, **2’**, or **2’’’** (200 mg/mL DMSO) | Monomer **1** (200 mL/mL) | Water | | |
| Non-cleavable TG (Monomer **2’** or **2’’**) | Single-round expansion | 2 parts (**2’** or **2’’**) | 1 part | 3 parts | | |
| Cleavable TG (Monomer **2’’’**) | 1st Gel for TG-based iExM | 2 parts (**2’’’**) | 1 part | 3 parts | | |
| Sodium polyacrylate/acrylamide gel (PAAG) | | | | | | |
| | | Acrylamide | Sodium acrylate | Crosslinker | PBS | NaCl |
| BAC-crosslinked expanding gel | 1st Gel for PAAG-based iExM | 2.6% | 8.6% | 0.2% BAC | 1x | 2M |
| BAC-crosslinked non-expanding gel | 2nd Gel | 10.4% | 0 | 0.2% BAC | 0 | 0 |
| DATD-crosslinked expanding gel | 3rd Gel | 2.6% | 8.6% | 0.5% DATD | 1x | 2M |
| DATD-crosslinked non-expanding gel | 4th Gel | 10.4% | 0 | 0.5% DATD | 0 | 0 |
| Bis-crosslinked expanding gel | 5th Gel | 2.6% | 8.6% | 0.15% bis | 1x | 2M |
